# Antifungal symbiotic peptide NCR044.1 exhibits unique structure and multi-faceted mechanisms of action that confer plant protection

**DOI:** 10.1101/2020.02.19.956318

**Authors:** Siva L. S. Velivelli, Kirk J. Czymmek, Hui Li, Jared B. Shaw, Garry W. Buchko, Dilip M. Shah

**Affiliations:** Donald Danforth Plant Science Center, St Louis, MO. 63132, USA; Advanced Bioimaging Laboratory, Donald Danforth Plant Science Center, St Louis, MO. 63132, USA; Earth and Biological Sciences Directorate, Pacific Northwest National Laboratory, Richland, WA 99354, USA; School of Molecular Biosciences, Washington State University, Pullman, WA 99164, USA

## Abstract

NCR044.1 is a 36-amino acid nodule-specific cysteine-rich antimicrobial peptide expressed in the developing nodules of *Medicago truncatula*. Here, we determined its unique NMR structure to be largely disordered, one four-residue α-helix and one three-residue anti-parallel β-sheet stabilized by two disulfide bonds, suggesting it is highly dynamic. NCR044.1 exhibited potent fungicidal activity against multiple plant fungal pathogens. It breached the fungal plasma membrane, bound to multiple phosphoinositides, and induced reactive oxygen species. Time-lapse confocal and super-resolution microscopy revealed strong fungal cell wall binding, penetration of the cell membrane at discrete foci, followed by gradual loss of turgor, and subsequent accumulation in the cytoplasm with elevated levels in nucleoli. Nucleolar localization of NCR044.1 was unique amongst plant antifungal peptides, suggesting its potential interaction with ribosomes and inhibition of translation. Spray-applied NCR044.1 significantly reduced gray mold disease symptoms caused by the fungal pathogen *Botrytis cinerea* in tomato plants and post-harvest products demonstrating its potential as a spray-on peptide-based biofungicide.

---

*Medicago truncatula*, a model legume of the inverted repeat-lacking clade (IRLC), forms an agriculturally important nitrogen-fixing endosymbiotic relationship with the Gram-negative bacterium *Sinorhizobium meliloti*^1^. This relationship results in the formation of indeterminate nodules on the plant’s roots whose continued peaceful existence depends on the mutual exchange of signals between the host and its natural symbiotic bacterial partner. After endocytosis into the cytoplasm of specialized nitrogen-fixing nodule cells, microsymbiont *S. meliloti* undergoes a remarkable irreversible differentiation process leading to the formation of enlarged polyploid bacteroides. This terminal differentiation process is orchestrated by 639 nodule-specific cysteine-rich (NCR) peptides expressed in successive spatio-temporal waves during nodule development^2–4^. Genetic studies have revealed that the absence of specific NCR peptides results in the breakdown of the symbiotic relationship and the cessation of nitrogen-fixation in *M. truncatula*^5–7^.

NCR peptides are directed to the plant-derived symbiosome containing bacteroids via the plant secretory pathway. Thus, each peptide is synthesized as a larger precursor peptide containing the amino-terminal endoplasmic reticulum targeting signal and the mature peptide. Mature NCR peptides are 30-50 amino acids in length but highly variable in their primary amino acid sequences and net charge. They are characterized by the presence of either four or six conserved cysteines presumably involved in formation of two- or three intramolecular disulfide bonds. NCR peptides are defensin-like since their predicted disulfide bonding patterns exhibit similarity with those of mammalian defensins. A subset of 639 NCR peptides is cationic with a net charge ranging from +3 to +11 and is also rich in hydrophobic residues^8^. Defensin-like NCR peptides exhibit potent antibacterial activity *in vitro* against symbiotic rhizobial bacteria and against various Gram-negative and Gram-positive bacteria^9–11^. For example, NCR247 has antibacterial activity against free-living *S. meliloti* at high concentrations but induces bacteroid-like features at low concentrations. A mode-of-action (MOA) study showed that NCR247 blocks cell division, induces cell elongation, and formed complexes with bacterial ribosomal proteins affecting protein synthesis and the bacterial proteome^12^. Since *M. truncatula* expresses several cationic NCR peptides, it is likely that they exhibit different modes of antibacterial action in different compartments of the nodule^13^.

In a recent study, 19 cationic NCR peptides were tested for their ability to inhibit the growth of a clinically relevant human fungal pathogen *Candida albicans*. Of these, nine with a net charge above +9 killed this fungal pathogen efficiently, showing inhibitory effects on both the yeast and hyphal forms at concentrations ranging from 1.5 to 10.5 μM^14^, demonstrating their potential for development as antifungal drugs^14^. The subcellular localization and intracellular targets of these peptides in fungal cells remain to be determined. There are several cationic NCR peptides with sequence heterogeneity expressed in the nodules of *M. truncatula* and other IRLC legumes and it is likely that at least some of them will have potent antifungal activity against plant fungal pathogens and will be unique in structure and MOA.

In this study, we showed that NCR044.1, a 36-amino acid peptide with a net charge of +9^14^, exhibited potent antifungal activity against plant fungal pathogens. However, its 3D structure was determined to be very different from other well characterized plant antifungal peptides such as defensins, thionins, lipid-transfer proteins, thaumatins, and heveins^15^. This is the first 3D structure of any NCR peptide and also the first antifungal plant peptide reported to target the nucleolus in fungal cells where it presumably binds to ribosomes and inhibits translation. NCR044.1 conferred strong resistance to *B. cinerea* when applied topically on lettuce leaves and rose petals. Young tomato and *Nicotiana benthamiana* plants sprayed with this peptide were also protected from the gray mold disease caused by this pathogen indicating the potential of NCR044.1 as a peptide-based fungicide. Our work illustrates the novel antimicrobial activity of NCR peptides and paves the way for development of these peptides as spray-on peptide-based biofungicides.

## Results

### Two NCR044 homologs are expressed in the nodules of *M. truncatula*

The *M. truncatula* genome contains genes encoding two NCR044 homologs, NCR044.1 (MtrunA17_Chr7g0216231) and NCR044.2 (MtrunA17_Chr7g0216241). The gene for each peptide encodes a signal sequence and a mature peptide. The mature peptides are each 36-amino acids in length and share 83% sequence identity (Fig. 1a). NCR044.1 has a net charge of +9 and 38% hydrophobic residues, whereas NCR044.2 has a net charge of +8 and 36% hydrophobic residues (Fig. 1b). Only two amino acid substitutions (I17P and R36S) account for the difference in net charge and hydrophobicity. The sequence of each mature peptide has a remarkably high concentration of cationic residues in the carboxy-terminal half of each molecule. The primary sequence of each peptide displays no significant homology with any of the peptide sequences in the Antimicrobial Peptide Database (http://aps.unmc.edu/AP/main.php)^16^.

**Figure 1.**
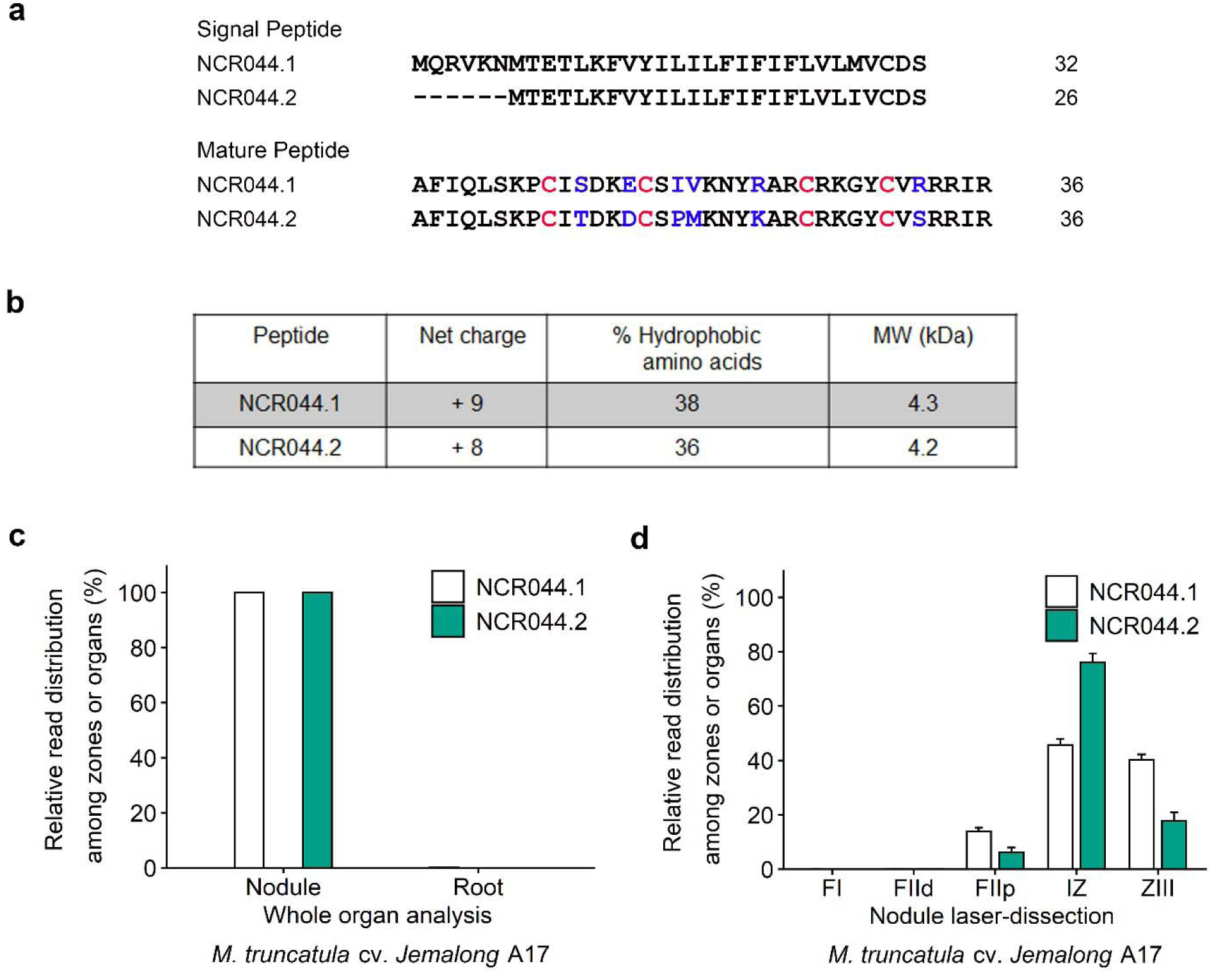
Amino acid sequences and properties of NCR044.1 and NCR044.2 and expression of their genes in root nodules of *Medicago truncatula* cv. *Jemalong* A17. **a.** Primary amino acid sequences of the signal and mature peptide sequences of NCR044.1 and NCR044.2. Mature peptide sequences are each 36 residues in length. Conserved cysteine residues are highlighted in red and six non-conserved residues are highlighted in blue. **b.** Net charge, hydrophobic amino acid content, and molecular weight of the mature NCR044.1 and NCR044.2 peptides. **c.** Gene expression analysis of NCR044.1 and NCR044.2 in root nodules and roots at 10 days post-infection (dpi) by *Sinorhizobium meliloti*. **d.** NCR044.1 and NCR044.2 transcript abundance in laser-dissected zones of mature nodules at 15 dpi. FI: nodule meristematic zone - Fraction I; FIId: infection zone – distal fraction; FIIp: early differentiation zone - proximal fraction; IZ: late differentiation zone - interzone II–III; ZIII: nitrogen-fixation zone. Web-based RNA-seq data are available at the Symbimics website (https://iant.toulouse.inra.fr/symbimics).

During development of indeterminate nodules, NCR044.1 and NCR044.2 transcripts are each highly expressed in the interzone II-III and the nitrogen-fixation zone ZIII and relatively lowly in the distal and proximal parts of the infection or differentiation zone (FIId and FIIp) (Fig. 1c, d)^17^. The biological function of each peptide during nodule development and nitrogen fixation remains to be determined.

### Expression of recombinant NCR044.1 in *Pichia pastoris* and mass spectral analysis of the purified peptide

In order to generate the NCR044.1 peptide with correctly formed disulfide bonds, the NCR044.1 gene was expressed in a heterologous *Pichia pastoris* expression system. The peptide secreted into the growth medium was successfully purified using cation exchange and C18 reverse phase HPLC. Mass spectral analysis revealed a molecular mass of 4311.28 Da for the purified product (Supplementary Fig. 1a, b) as expected for disulfide-linked NCR044.1.

### Disulfide bonding pattern in NCR044.1

NCR044.1 contains four cysteine residues (C9, C15, C25, C31) with three possible patterns of intramolecular disulfide bond formation and multiple patterns of intermolecular disulfide formation. Since the intra- and intermolecular pattern of disulfide bond formation will significantly affect the peptide’s structure, and likely its function, it is important that oxidation is primarily intramolecular and the pattern of oxidation in NCR044.1 is homogeneous. NMR measurement of the peptide’s rotational correlation time (τ_c_), 3.6 ± 0.5 ns, was consistent for a ∼4.0 kDa peptide^18^ indicating it was primarily monomeric in solution with no detectable intermolecular disulfide bond formation.

Fig. 2a and 2b show the assigned ^1^H-^15^N HSQC spectra for the oxidized and reduced forms of NCR044.1. For oxidized NCR044.1 (Fig. 2a), the wide chemical shift dispersion of the amide resonances in both the proton and nitrogen dimensions are characteristic features of a structured protein^19^. A single set of cross peaks that could all be assigned suggests one dominant species is present with a homogeneous intramolecular disulfide bond pattern. Disulfide bond formation in NCR044.1 was unambiguously verified from the NMR chemical shifts of the β-carbon of cysteine residues. Generally, the cysteine ^13^C^β^ chemical shift in the oxidized state is >35 ppm and decreases to <32 ppm in the reduced state^20^. In the absence of tris(2-carboxyethyl)phosphine (TCEP) or other reducing agents the cysteine ^13^C^β^ chemical shifts for NCR044.1 were 38.0 ppm or greater, as indicated in Supplementary Table 1, clearly indicating that cysteine residues were oxidized and incorporated in disulfide bonds. These ^13^C^β^ chemical shifts decreased to 28 ppm or less in the presence of 5 mM TCEP, a significant change indicating that disulfide bonds were ablated.

**Figure 2.**
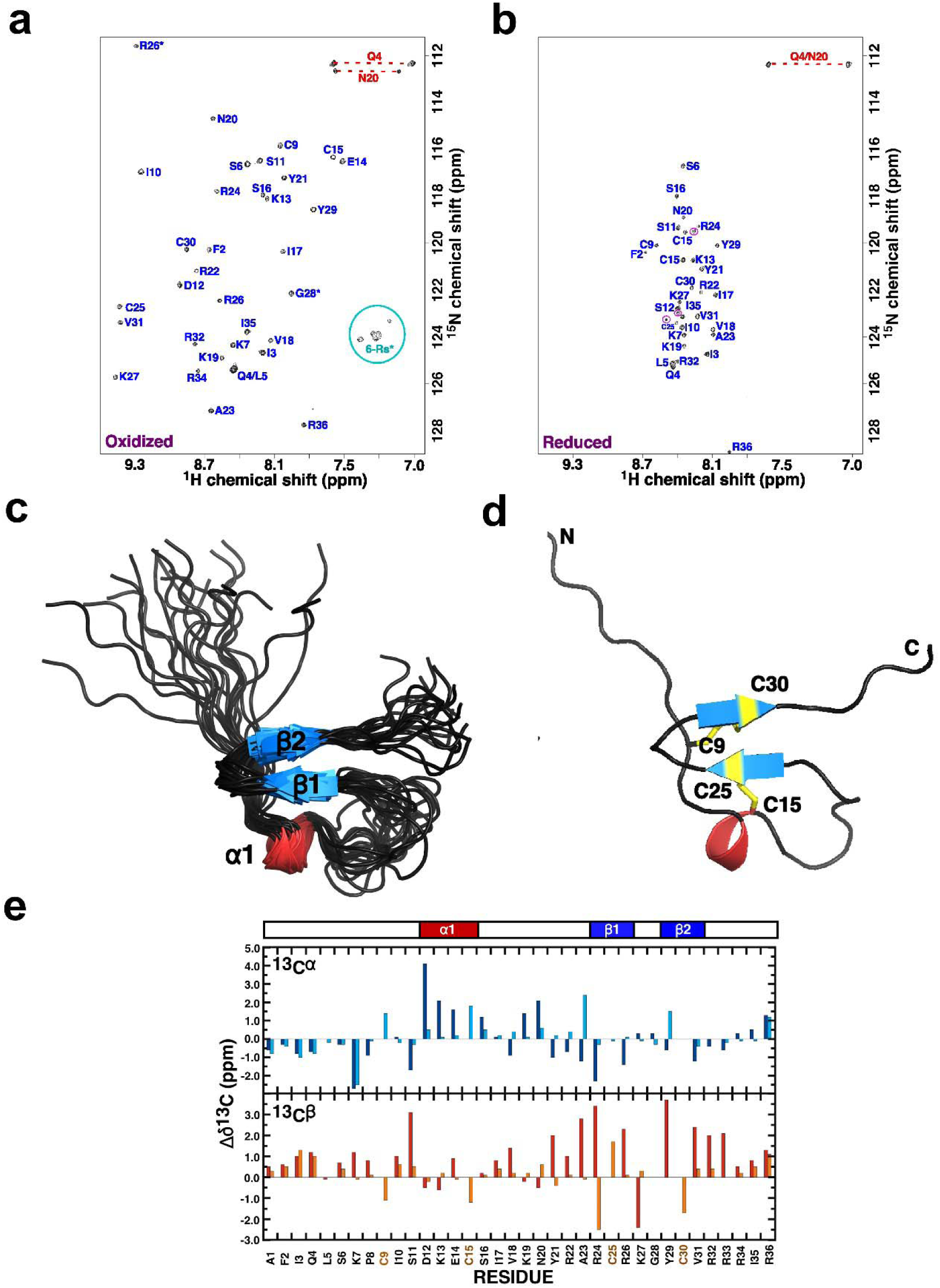
Solution structure of NCR044.1. **a-b**. Assigned ^1^H-^15^N HSQC spectra for NCR044.1 (0.7 mM) collected in the oxidized (disulfide) (**a**) and reduced (dithiol) (**b**) states. Spectra collected at 20 °C in 20 mM sodium acetate, 50 mM NaCl, pH 5.3 at a ^1^H resonance frequency of 600 MHz. Amide side chain resonance pairs are connected by a red dashed line, guanidium protons of the six arginine residues are in the cyan circle (not visible in the reduced state), and folded resonances are identified with an asterisk. The three cross peaks circled in magenta in reduced NCR044.1 are resonances for R26, R32, or R33 that could not be unambiguously assigned. **c.** Cartoon representation of the backbone superposition of the ordered regions in the ensemble of 19 structures calculated for oxidized NCR044.1 (6U6G). Beta-strands are colored blue and the α-helix is colored red. **d.** Cartoon representation of the NCR044.1 structure closest to the average with a stick representation of the four cysteine side chains highlighted in yellow. **e.** Analysis of the side chain ^13^C chemical shifts in the reduced (dithiol) and oxidized (disulfide) states. The observed ^13^C^α^ and ^13^C^β^ chemical shift deviations from random coil values for oxidized and reduced NCR044.1 where Δ^δ^13^C^ = δ_Observed_ – δ_Randon coil_. The random coil carbon values were taken from CNS (cns_solve_1.1) with no calculations for the oxidized cysteine residues. Blue and red = oxidized, cyan and orange = reduced. On top of the graph is a schematic illustration of the elements of secondary structure observed in the NMR-derived structure with the α-helix colored red and β-strands colored blue.

While the chemical shift data indicated disulfide bonds were present, it was not possible to use preliminary structure calculations to deduce unambiguously the disulfide bond pairs from ^1^H-^1^H NOEs due to the consequences of proximal γ-sulphur atoms. Instead, the disulfide bond pattern was deduced from the analysis of mass spectral data for trypsin-digested, unlabeled peptide in the reduced and oxidized states. This data showed that disulfide linkages were C9-C30 and C15-C25. Note that the disulfide bonds are essential for the stability of the peptide’s tertiary structure^21^ as illustrated in the ^1^H-^15^N HSQC spectrum for reduced NCR044.1 in Fig. 2b. The range of chemical shift dispersion of amide resonances was greatly compressed relative to the oxidized NCR044.1 (Fig. 2b), a characteristic feature of disordered proteins^19^.

### Solution NMR structure of NCR044.1

Fig. 2c is a superposition of the final ensemble of structures calculated for NCR044.1 (PDB: 6U6G) with a single cartoon representation of this ensemble shown in Fig. 2d. The most striking feature is the paucity of canonical elements of secondary structure in the peptide. NCR044.1 contains one short (A23-R25, G28-C30) anti-parallel β-sheet and a “whif” (S11-E14) of an α-helix. The rest of the peptide was largely disordered and dynamic, even with the tethering afforded by the two disulfide bonds. A search for structures in the PDB similar to NCR044.1 with the DALI server^22^ generated no hits, suggesting this structure was new to the PDB. The disorder in the structure was reflected in the paucity of ^1^H-^1^H NOEs observed in the NOESY NMR data. As shown in Supplementary Table 2, most of the NOEs were intra-residue with only 12 long range NOEs observed. The prevalent disorder was also reflected in the analysis of the ^13^C^α^ and ^13^C^β^ chemical shifts. Deviations of ^13^C^α^ and ^13^C^β^ chemical shifts from random coil values are predictive of canonical α-helical (positive Δδ^13^C^α^, negative Δδ^13^C^β^) and β-strand (negative Δδ^13α^, positive Δδ^13^C^β^) secondary structure^21,23^. These deviations from random coil values were plotted for oxidized and reduced NCR044.1 in Fig. 2e and showed modest correlation for their respective short elements of the secondary structure identified in the NMR structure calculations for the oxidized peptide. Upon reducing the disulfide bonds in the presence of 5 mM TCEP, aside from the N-terminal region (A1 – I10) which was largely disordered in the oxidized state to begin with, these modest correlations disappear with most of the chemical shift values not significantly differing from random coils values. These observations suggest the disulfide bonds play a significant role in stabilizing the structure of the oxidized peptide and in their absence the peptide was largely disordered with little transient structure.

### NCR044.1 exhibits potent antifungal activity against plant fungal pathogens

Chemically synthesized NCR044.1 was previously reported to exhibit fungicidal activity against the human fungal pathogen *C. albicans* at low micromolar concentrations^8^. Here, we determined the antifungal activity for recombinant NCR044.1 against a panel of closely related plant fungal pathogens, *Fusarium* spp. and *B. cinerea*, using a spectrophotometric assay. NCR044.1 inhibited the growth of *B. cinerea* at low micromolar concentration with an IC_50_ value of 1.55 ± 0.21 μM (Fig. 3a). NCR044.1 also inhibited the growth of *F. graminearum* and *F. virguliforme* with IC_50_ values of 1.93 ± 0.23 and 1.68 ± 0.20 μM, respectively. *F. oxysporum* was the most sensitive to NCR044.1 with an IC_50_ value of 0.52 ± 0.01 μM (Fig. 3a). The resazurin cell viability assay revealed that *F. oxysporum* cells were killed at a concentration of 1.5 μM, whereas other fungi, including *B. cinerea*, were killed at a concentration of 3 μM (Fig. 3b and 3c).

**Figure 3.**
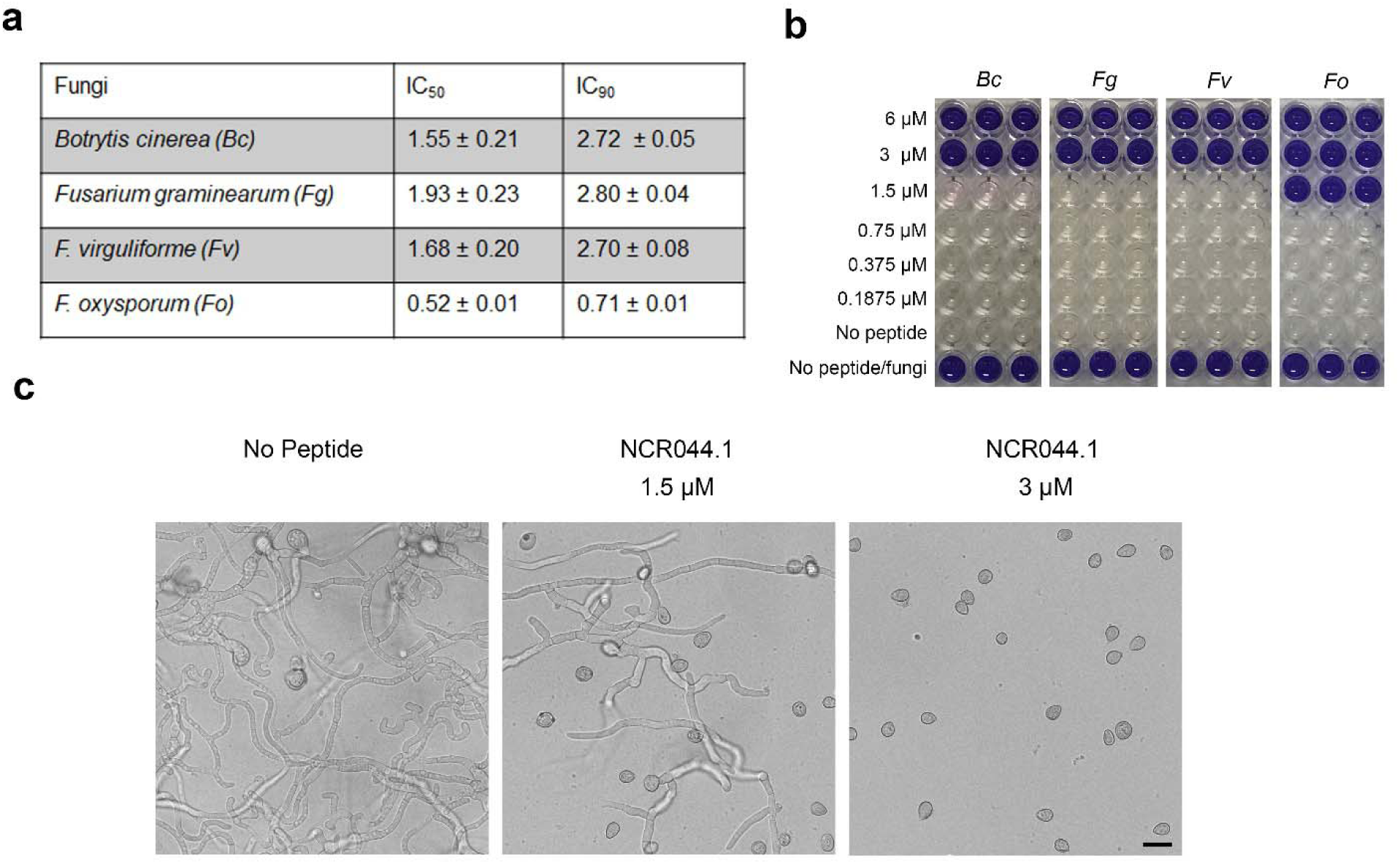
Antifungal activity of NCR044.1 against *B. cinerea* and *Fusarium* spp. **a.** IC_50_ and IC_90_ values of NCR044.1 for each pathogen are shown. Data are means ± SEM of three independent biological replications (n = 3). **b.** Results of the fungal cell viability assay using resazurin, a metabolic indicator of living cells. A change from blue to pink/colorless signals resazurin reduction and indicates metabolically active fungal cells. In the presence of 3 or 6 μM NCR044.1, fungal cells lost their metabolic capacity and did not reduce resazurin **c.** Representative microscopic images showing the inhibition of *B. cinerea* growth 24 - 48 h after treatment with 1.5 or 3 μM of NCR044.1 (right). *B. cinerea* without peptide added served as a negative control (left). (Scale bar: 20 μm).

### NCR044.1 disrupts the plasma membrane of *B. cinerea*

Antifungal peptides are known to interact with phospholipid bilayers and disrupt the plasma membrane^15^. We postulated that the antifungal activity of NCR044.1 occurred via membrane permeabilization. Using a SYTOX^TM^ Green (SG) assay, a non-fluorescent intercalating dye that permeates fungal cells when their plasma membrane integrity is compromised, we tested the ability of NCR044.1 to cross the plasma membrane of *B. cinerea*.

The uptake of SG in *B. cinerea* conidia and germlings treated with 3 μM NCR044.1 for 15 min was first monitored using confocal microscopy. The nuclei of both conidia and germlings were stained with SG, suggesting compromised cellular membrane integrity induced by NCR044.1 (Fig. 4a-d). The kinetics of membrane permeabilization was determined in *B. cinerea* germlings treated with various concentrations of NCR044.1 every 30 min for up to 3 h. In NCR044.1 treated fungal cells, permeabilization of the plasma membrane was observed within 30 min and reached its maximal level at 120 min. The rate of membrane permeabilization increased with increasing concentrations of NCR044.1 as shown in Fig. 4i. To test if membrane disruption was sufficient for antifungal activity of NCR044.1 in ungerminated *B. cinerea* spores, we incubated spores in 3 μM NCR044.1 for 1 h, washed off free peptide, and then allowed the spores to germinate for 24 h at room temperature in peptide-free growth medium. Our data showed that *B. cinerea* spores were able to resume their growth (Fig. 4j-l) in peptide-free growth medium suggesting membrane permeabilization alone is insufficient to induce death of fungal cells by the peptide.

**Figure 4.**
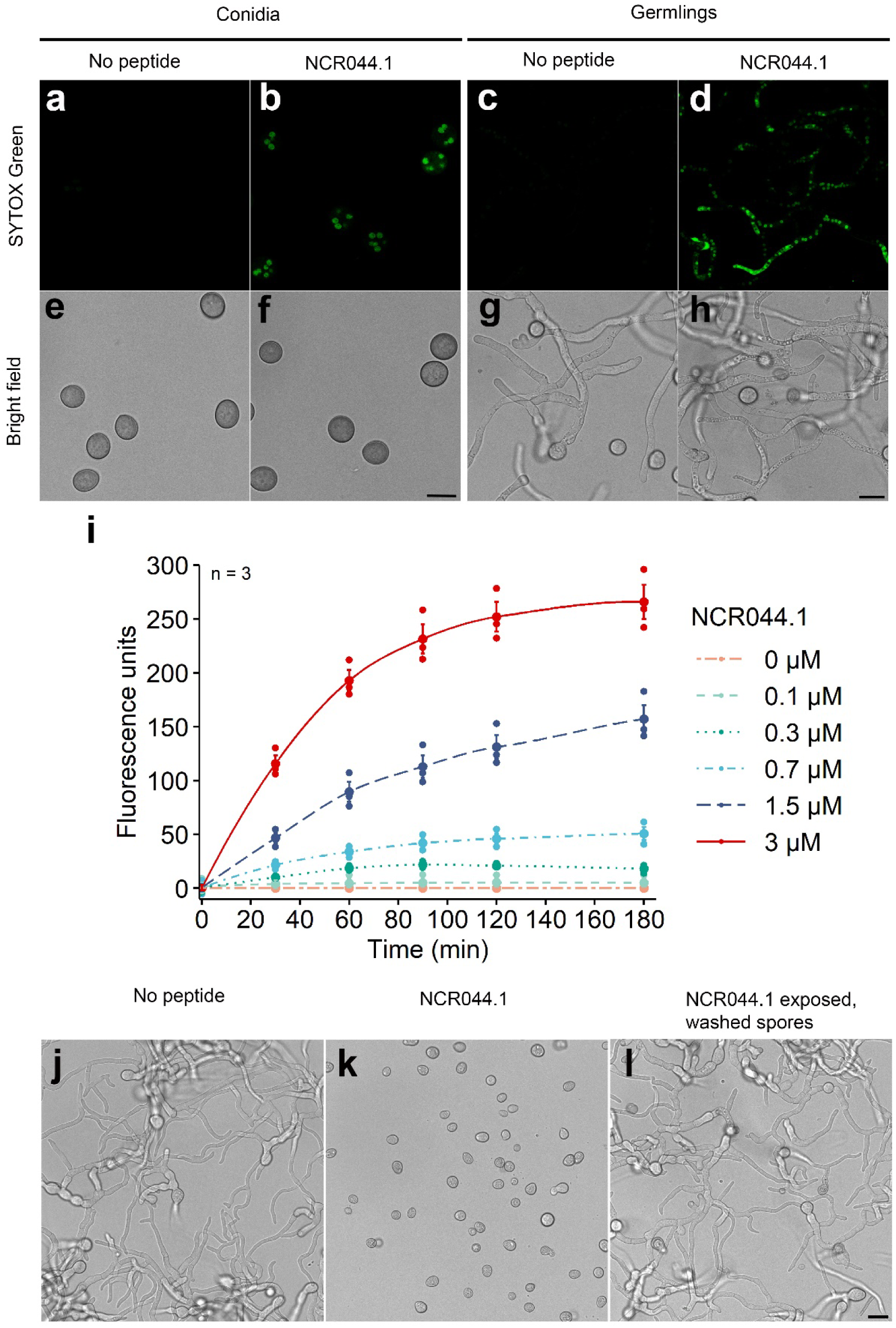
Membrane permeabilization activity of NCR044.1. **a-d.** Confocal microscopy images**, e-h**. Corresponding bright field images (e-h) of SYTOX^TM^ Green (SG) uptake in *B. cinerea* conidia and germlings treated with 3 μM NCR044.1 for 15 min. (Scale bar: 10 μm). SG (green) labeled nuclei in conidia and germlings treated with NCR044.1 indicate increased plasma membrane permeability. **i.** Real time-quantification of cell membrane permeability in *B. cinerea* germlings treated with various concentrations of NCR044.1. Membrane permeabilization increased with increasing concentrations of peptide. Data are means (large dots) ± SEM of three biological replications (n = 3, small dots). **j.** Representative microscopic images of *B. cinerea* spore germination without peptide (left). **k.** *B. cinerea* growth inhibition in the presence of 3 μM NCR044.1 (middle). **l.** The germination of *B. cinerea* spores 24 h after removal of 3 μM NCR044.1 after treatment for 1 h (right). (Scale bar: 10 μm). Fungal growth after removal of the peptide indicates that membrane permeabilization is insufficient for fungal cell killing.

### NCR044.1 elicits rapid production of ROS in *B. cinerea* germlings, but not in conidia

Antimicrobial peptides with different mode of action are known to induce ROS and cause cell death through apoptosis or necrosis-like processes^24, 25^. To determine if NCR044.1 elicited production of ROS, we challenged the conidia and germlings of *B. cinerea* with 3 μM of NCR044.1 for 2 h in the presence of the dye 2′,7′-dichlorodihydrofluorescein diacetate (H_2_DCF-DA). Upon intracellular oxidation by ROS, H_2_DCF-DA is converted into a highly fluorescent 2’,7’-dichlorofluorescein (DCF) that can be monitored using confocal microscopy. ROS production was observed in germlings, but not in conidia (Supplementary Fig. 2a-d). Real time-quantification of ROS production was measured spectrophotometrically every 30 min for up to 2 h with untreated germlings and germlings treated with 3 μM NCR044.1. Rapid induction of ROS fluorescence within 30 min was observed and the fluorescence intensity increased in a time- and dose-dependent manner. The highest fluorescence intensity was observed at 3 μM suggesting the elicitation of oxidative stress by ROS in germlings following NCR044.1 treatment (Supplementary Fig. 2i).

### NCR044.1 binds to multiple membrane phospholipids

Several plant defensins have been shown to recruit plasma membrane-resident phospholipids as part of their MOA^26, 27^. Using the protein-lipid overlay assay, we assessed the ability of NCR044.1 to bind different bioactive membrane phospholipids. NCR044.1 strongly bound to phosphatidylinositol 3,5-bisphosphate (PI(3,5)P_2_). It also bound weakly to multiple other phosphoinositides and to phosphatidic acid (PA) (Fig. 5a, b). To further investigate the phospholipid-binding specificity of NCR044.1, a polyPIPosome assay was performed with polymerized-liposomes. As shown in Fig. 5c, NCR044.1 bound strongly to liposomes that contained PC:PE:PI(4)P and PC:PE:PI(3,5)P_2_ and weakly to PC:PE containing liposomes.

**Figure 5.**
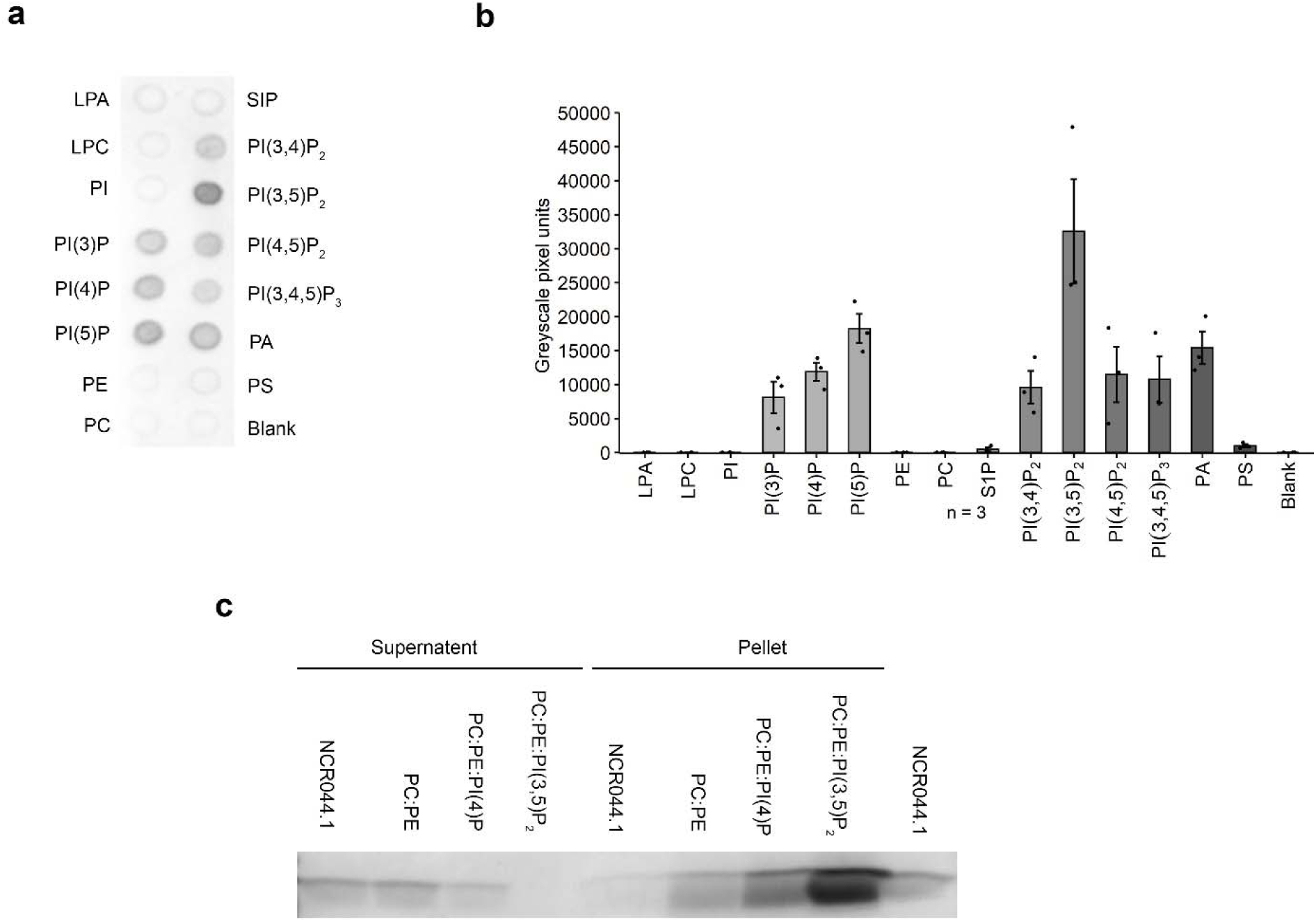
Phospholipid binding of NCR044.1. **a.** PIP strip showing strong binding of NCR044.1 to phosphatidylinositol diphosphate PI(3,5)P_2_ and weak binding to multiple phospholipids, including phosphatidylinositol monophosphates PI(3)P, PI(4)P and PI(5)P, phosphatidylinositol di/tri-phosphates PI(3,4)P_2_, PI(4,5)P_2_, PI(3,4,5)P_3_, and phosphatidic acid (PA). **b.** Densitometry analysis of PIP strip probed with NCR044.1. Data are means ± SEM of three independent biological replications (n = 3, small dots). **c.** PolyPIPosome binding assay showing strong binding of NCR044.1 to PI(4)P and PI(3,5)P_2_ and weak binding to PC:PE.

### NCR044.1 interacts with PI3P to form a large molecular weight complex

On the basis of the above protein-phospholipid overlay assay, we conducted an NMR chemical shift perturbation study with ^15^N-labeled oxidized NCR044.1 and PI(3)P to further corroborate phospholipid binding, and ideally, identify the lipid binding surface of NCR044.1. This experiment is based on the premise that backbone amide protons are sensitive to their molecular environment, and consequently, it is possible to identify the surface of ligand binding to a polypeptide by following amide chemical shift (or intensity) perturbations in ^1^H-^15^N HSQC spectra following the titration of ligand into the sample^28, 29^. For NCR044.1, no chemical shift perturbations were observed over the titration range, but the intensity of the amide cross peaks was observed to decrease starting at a PI(3)P:NCR044.1 molar ratio of 9.4:1. As shown in Supplementary Fig. 3a, there were remnants of only 12 amide cross peaks at a PI(3)P:NCR044.1 molar ratio of 18.8:1. These intensity perturbations in the ^1^H-^15^N HSQC spectra of NCR044.1 showed that the peptide was interacting with PI(3)P^30^. This was corroborated by overall rotational correlation time (τ_c_) measurements of the peptide at each titration point that increased from 3.6 ± 0.5 ns in the absence of PI(3)P to 8.7 ± 2.5 ns at a PI(3)P:NCR044.1 molar ratio of 18.8:1. The latter τ_c_ value corresponded to a molecular weight in the 15 kDa range^18^. Because amide cross peaks should be readily detectable in the ^1^H-^15^N HSQC spectrum of an ∼15 kDa species, potential explanations for the disappearing amide resonances include (i) heterogeneous binding to the phospholipid that generates multiple different chemical environments for the peptide with populations that cannot be detected and (ii) dynamic interactions with a large molecular complex formed by PI(3)P in solution. To the best of our knowledge, the solution properties of PI(3)P have not been characterized at high concentrations. However, given the aliphatic tails on one end of the molecule and the hydrophilic inositol head-group on the other end, it is possible that PI(3)P forms high molecular weight micellar structures at high lipid concentrations. Interactions between PI(3)P and NCR044.1 are likely electrostatic in nature. The hydrophilic head-group of PI(3)P contains two negatively charged phosphate groups. As shown in Supplementary Fig. 3b, the surface of NCR044.1 is dominated by a positively charged surface that covers the majority of the peptide, providing multiple sites for binding to the hydrophilic head-group of PI(3)P.

### Exogenous NCR044.1 is translocated into the cells of *B. cinerea*

We used DyLight550-labeled NCR044.1 to examine its intracellular translocation into the *B. cinerea* conidia and germling cells. In conidia incubated with 3 μM DyLight550-NCR044.1 for 16 h, the peptide first accumulated on the cell surface, and was subsequently internalized and diffused throughout the cytoplasm (Fig. 6a, b). We quantified the percentage of conidia cells that internalized 1.5 and 3 μM DyLight550-labeled NCR044.1 after overnight incubation and observed a statistically significant difference (*p* = 0.0057, *t* = −5.4, df = 4, unpaired Student’s *t* test) (Fig. 6c). In conidial cells treated with 1.5 μM peptide, only 18.8% of the cells internalized peptide into the cytosol. In contrast, this value rose to 47.5% with 3 μM peptide. The membrane selective dye FM4-64 was subsequently used to determine if DyLight550-labeled NCR044.1 co-localized with cellular membranes in *B. cinerea* germlings. In spore heads and germlings, NCR044.1 first bound to the cell wall (Fig. 6d-f, arrows) and cell membrane (Fig. 6d-f, arrowheads). NCR044.1 co-localized with FM4-64 at bright foci at the germ tube tip and small foci intermittently along the cell periphery of *B. cinerea* (Fig. 6 g-i, asterisks). This suggested that this peptide likely interacted with plasma membrane resident phospholipids which invaginated and/or endocytosed to enter fungal cells. Additionally, we observed that NCR044.1 was distributed diffusely in the cytoplasm and concentrated within the nucleus (Fig. 6d-i, N), whereas FM4-64 showed clear cytoplasmic membrane profiles and was excluded from the nucleus, suggesting that NCR044.1 localized at other non-membrane intracellular targets.

**Figure 6.**
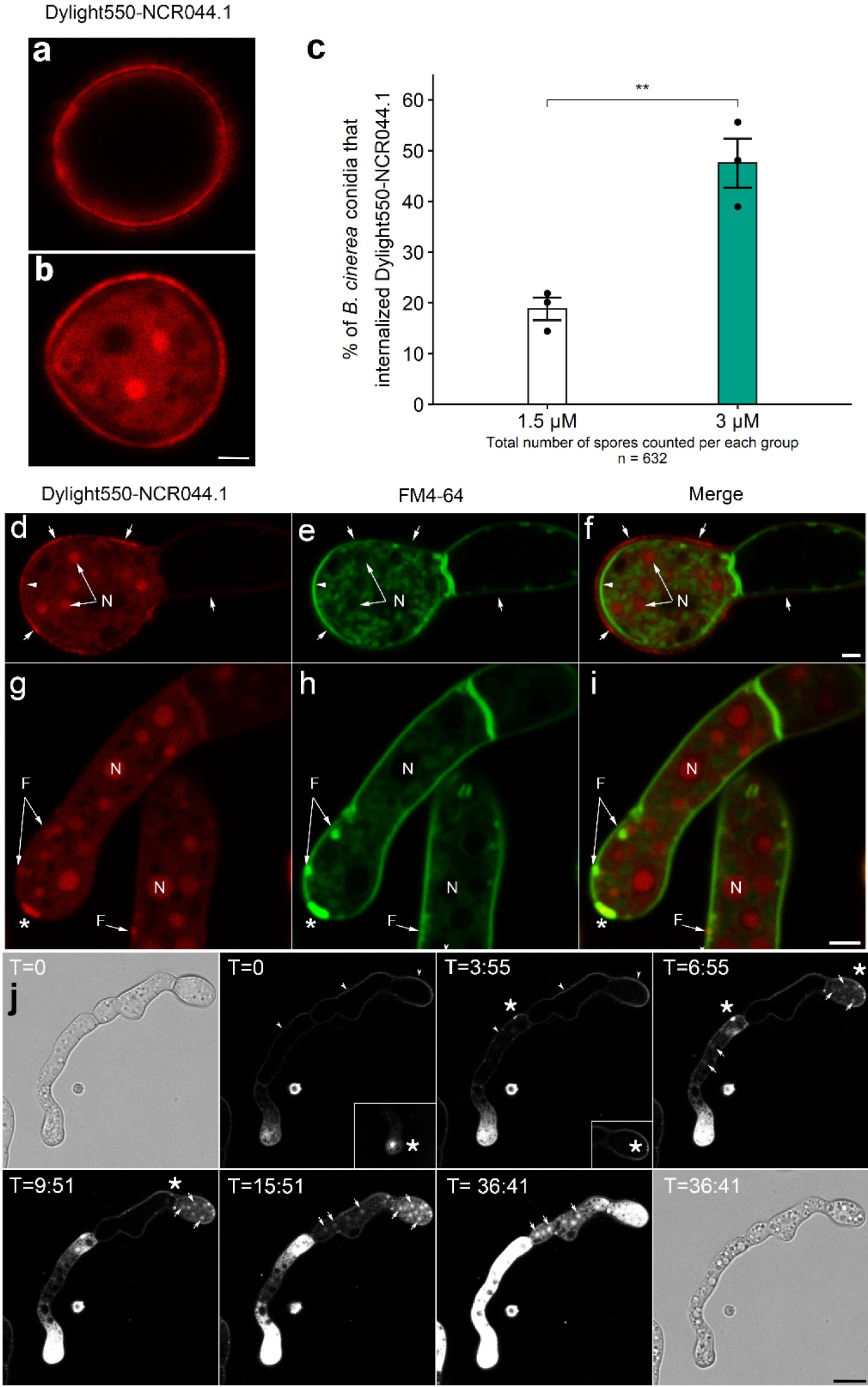
Translocation of exogenous NCR044.1 into *B. cinerea* cells. **a-b.** Confocal microscopy images of *B. cinerea* conidia showing the uptake of DyLight550-NCR044.1. NCR044.1 first accumulated on the cell surface and was internalized inside the *B. cinerea* conidia after overnight incubation (Scale bar: 2 μm). **c.** Percentage of *B. cinerea* conidia that internalized DyLight550-NCR044.1 following overnight incubation with 1.5 and 3 μM labeled peptide. Data are means ± SEM of three independent biological replications (n = 3, small dots). Asterisks denote significant differences (**P<0.01, unpaired Student’s t test). **d-i**. Confocal microscopy images of *B. cinerea* spores and germlings showed the internalization and co-localization of 3 μM Dylight550-NCR044.1 (red) with membrane-selective dye FM4-64 (green). DyLight550-NCR044.1 bound to the cell wall (arrows) and cell membranes (arrow heads). DyLight550-NCR044.1 co-localized with FM4-64 and bright focal accumulations at the germ tube tip (asterisks) and intermittently adjacent to cell walls (F) of *B. cinerea*. DyLight550-NCR044.1 localization to nuclear region (N) (Scale bar: 2 μm). **j.** Time-lapse confocal microscopy images of *B. cinerea* germlings showed internalization of Dylight550-NCR044.1 (white) over ∼ 40 min. Peptide first accumulated along cell wall (T = 0), and then entered inside the *B. cinerea* at foci (asterisks) at the germ tube tip or along the germ tube and/or conidial walls (T = 0 through T = 9:51 minutes). Peptide was diffusely localized into the cytoplasm and a strong signal was observed throughout the nuclear region of spore heads and germlings (T = 9:51 through 36:41 minutes). Bright field images at T = 0 and T = 36:41 show loss of turgor and cytoplasmic vacuolization (Scale bar: 10 μm).

Next, we applied time-lapse confocal microscopy to monitor the uptake of 3 μM DyLight550-labeled NCR044.1 in germlings. The peptide initially was observed associated with the germling cell wall (Fig. 6j, T=0) and then migrated towards foci at the germ tube tip (Fig. 6d, T=0, asterisks) or the germ tube and/or conidial walls (Fig. 6j, T=0 through T=9:51, asterisks). These foci appeared to be the sites of cytoplasmic entry and coincided with a gradual loss of turgor (Compare Fig 6J T=0 with T=40). Within 15 min, the peptide, in general, was diffusely localized in the cytoplasm of the germlings and tended to associate with bright spherical structures consistent with nuclei. Often, entry into the cell was observed as a gradient across the cell dependent on the origin of plasma membrane breach (Fig. 6j).

### Super-resolution microscopy reveals nucleolus as a cellular target of NCR044.1 in *B. cinerea*

We applied lattice structured illumination microscopy to clearly identify the subcellular localization of DyLight550-labeled NCR044.1 in *B. cinerea* at high-resolution. Since we determined that DyLight550-labeled NCR044.1 was preferentially localized to the nucleus of *B. cinerea* (Fig. 7a, b), we applied the DNA staining dye DAPI to see if the peptide bound to the nuclear DNA. As shown in Fig. 7c, DyLight550-NCR044.1 did not co-localize with DAPI in the nuclei of the germlings cells, and indeed, appeared to be explicitly excluded from nuclear DNA. We then used the nucleolus-specific RNA staining dye, Nucleolus Bright Green (NBG), to monitor the co-localization with the peptide. We observed that DyLight550-NCR044.1 strongly co-localized with NBG (Fig. 7d) suggesting potential interaction with ribosomes. Thus, in *B. cinerea*, NCR044.1 appears to be a nucleolus-targeting peptide with potential to form a complex with ribosomes and impair their function.

**Figure 7.**
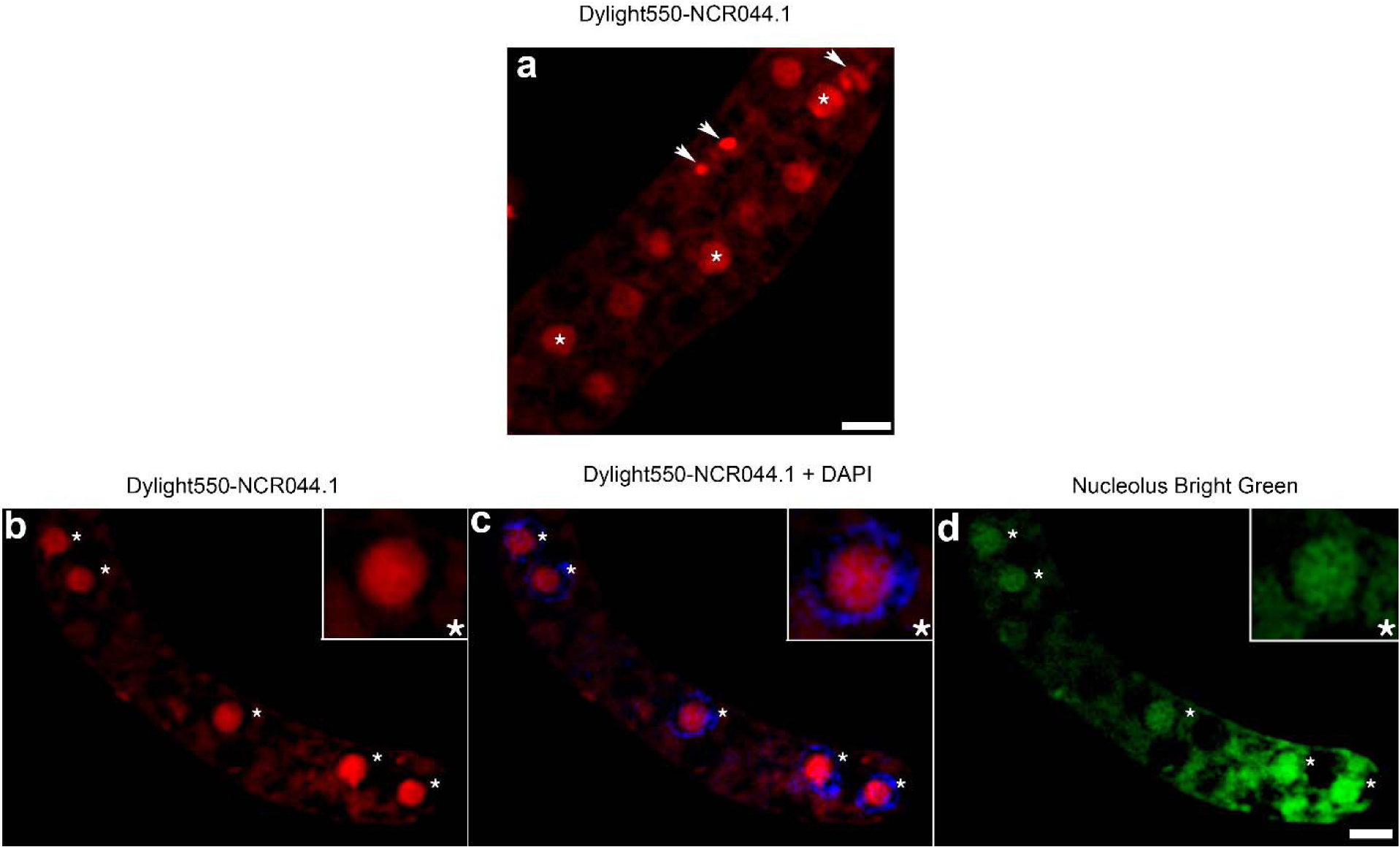
Subcellular localization of fluorescently labeled DyLight550-NCR044.1 in *B. cinerea* germlings. **a.** Super-resolution structured illumination microscopy (SR-SIM) images of *B. cinerea* germlings showed the internalization of DyLight550-NCR044.1 (red). DyLight550-NCR044.1 was targeted at elevated levels to the nucleus in *B. cinerea* germlings (asterisks). **b-d.** DyLight550-NCR044.1 (**b**) did not co-localize with the DNA-specific DAPI (asterisks) stain (**c**), but, co-localized with rRNA specific Nucleolus Bright Green stain (**d**) suggesting that NCR044.1 specifically targeted the nucleolus. Images were taken after 1 h of exposure to 3 μM DyLight550-NCR044.1 (Scale bar: 2 μm).

### Fluorescence Recovery After Photobleaching (FRAP) experiments reveal that NCR044.1 has strong binding affinity to *B. cinerea* conidia cell walls and increased affinity to the nucleolus in the cell

In order to understand the mobility of DyLight550-NCR044.1 to specific subcellular domains in *B. cinerea*, we performed a series of FRAP experiments (Fig. 8a-e) quantifying the half time of recovery (τ_½_) of the mobile fraction and percent immobile fraction. We targeted four distinct regions; the conidial spore wall, germ tube cell wall, cytoplasm, and nucleoli, using identical FRAP bleach/recovery acquisition and analysis conditions. Statistically significant differences (K-W_statistic_ = 59.79, df = 3, *p* < 0.0001) were observed between specific cellular structures and among the immobile fractions of spore walls versus germ tube walls, cytoplasm, and nucleoli (pairwise Wilcoxson test, corrected *p* < 0.0001) (Fig. 8b). Our data showed that the fungal cell walls had the highest affinity to DyLight550-NCR044.1 with an immobile fraction of 76.7 ± 1.3% (spores) and 24.9 ± 3.3% (germ tubes), while the nucleoli and cytoplasm had greatly reduced immobile fractions of 15.4 ± 1.9 and 11.7 ± 1.5%, respectively (Fig. 8c). Additionally, the τ_½_ of the mobile fractions revealed a statistically significant difference (K-W_statistic_ = 27.81, df = 3, *p* < 0.0001) between nucleoli versus cytoplasm (pairwise Wilcoxson test, corrected *p* < 0.01) and nucleoli versus cell wall (corrected *p* < 0.01) (Fig. 8d). Spore walls and nucleoli had the slowest mobility with a τ_1/2_ = 36.0 ± 2.4 and 31.4 ± 2.4 seconds, respectively, while the cytoplasm and germ tube cell walls had the fastest mobility with a τ_1/2_ = 22.0 ± 1.7 and 20.5 ± 3.4 seconds, respectively (Fig. 8e). Taken together, this data suggested that strong binding of the spore cell wall by NCR044.1 might serve to anchor it in close proximity to the plasma membrane and then upon germination, the mobility of NCR044.1 increased, allowing it to breach the cell membrane and migrate to cytoplasmic targets with preferential affinity to the nucleoli.

**Figure 8.**
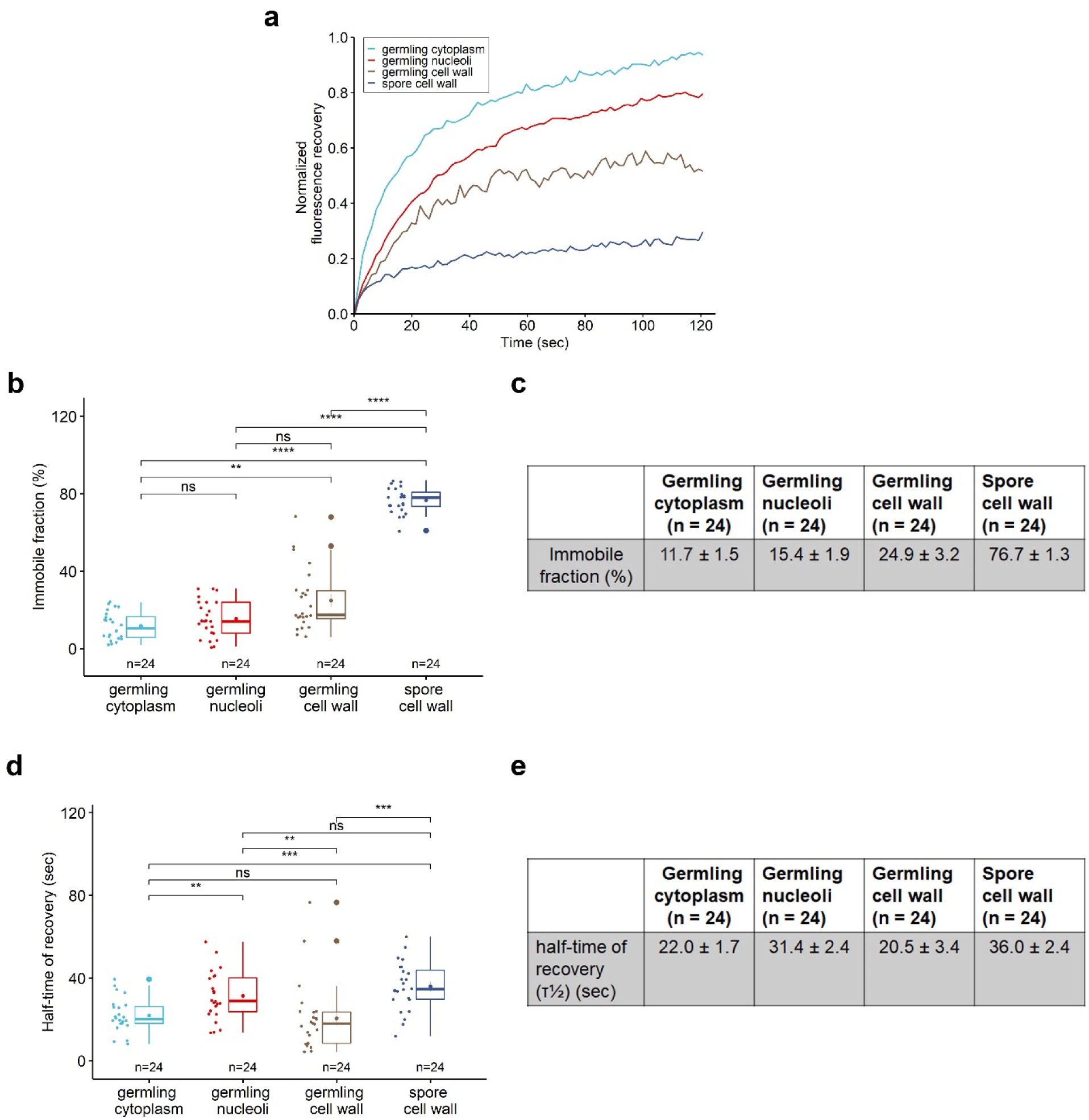
Fluorescence Recovery After Photobleaching. **a.** Representative normalized fluorescence recovery curves of DyLight550-NCR044.1 showed relative differences in mobility of key subcellular compartments in *B. cinerea*. DyLight550-NCR044.1 recovery in the germling cytoplasm (light blue) had the greatest overall mobility compared with germling nucleoli (red), germling cell walls (brown) and spore cell walls (blue). Spore cell walls exhibited the least mobility (dark blue) of all measured compartments. Recovery data derived from **a** was normalized to 1 and the x and y-axis was set to the origin “0”. **b.** The immobile fraction (%) data are represented using a boxplot and tabulation of the average values (**c).** The immobile fraction (representing tightly bound peptide) was relatively low for the cytoplasm (11.7 ± 1.5 %), nucleoli (15.4 ± 1.9 %), and germling cell walls (24.9 ± 3.2 %) but significantly greater for spore cell walls (76.7 ± 1.3 %). **d**. Half-time of recovery (τ½) data are represented using a boxplot and tabulation of the average values (**e)**. NCR044.1 mobile fractions showed relatively quick recovery (τ½ = 22.0 ± 1.7 s) in the germling cytoplasm and germling cell walls cell wall (τ½ = 20.5 ± 3.4 sec) while nucleoli (τ½ = 31.4 ± 2.4 s) and spore cell walls (τ½ = 36.0 ± 2.4 s) exhibited reduced mobility suggesting stronger interactions with these structures. Each colored boxplot represents the 25^th^ and 75^th^ percentiles (box) and the whisker represents 1.5 times the interquartile range (IQR) from the 25th and 75th percentiles. The horizontal line in the box plot represents the median with the mean ± SEM denoted by a dot. Outliers are indicated by large dots outside 1.5*IQR above the first and third quartile. The individual measurements of mobile and immobile fractions are indicated on the left of each boxplot and represent the variances of the samples. The number (n = 24) of different subcellular compartments in *B. cinerea* analyzed are indicated below each group in the boxplot and are from at least three independent experiments. Asterisks represent significant differences between different subcellular compartments (*P<0.05, **P<0.01 ***P<0.001, ****≤0.0001, pairwise Wilcoxson test with Holm correction, Kruskal–Wallis for multiple groups).

### NCR044.1 confers resistance to *B. cinerea* in lettuce leaves and rose petals

NCR044.1 was tested for its ability to reduce symptoms of gray mold disease *in planta.* Detached lettuce leaves were drop-inoculated with the conidia of *B. cinerea* in the presence of different concentrations of the peptide. Significant differences in disease severity were found between plants treated with peptide and control plants without peptide treatment (Kruskal-Wallis_statistic_ = 31.15, df = 3, *p* < 0.0001). In particular, lettuce leaves inoculated with the pathogen in the presence of 6 and 12 μM NCR044.1 displayed significantly smaller lesions 48 h post-inoculation compared to control leaves sprayed with pathogen only (pairwise Wilcoxson test, corrected *p* < 0.0001) (Supplementary Fig. 5a).

NCR044.1 was also tested for its ability to confer resistance to *B. cinerea* in a rose petal infection assay. NCR044.1 significantly reduced virulence of the pathogen on rose petals as compared with control petals without peptide treatment (K-W_statistic_ = 76.36, df = 3, *p* < 0.0001). Almost complete suppression of disease symptoms was observed at a concentration of 1.5 μM peptide compared to rose petals sprayed with pathogen only (pairwise Wilcoxson test, corrected *p* < 0.0001) (Supplementary Fig. 5b).

### Spray-applied NCR044.1 confers gray mold resistance in *N. benthamiana* and tomato plants and is not internalized by plant cells

We tested the potential of NCR044.1 for use as a peptide fungicide for controlling gray mold disease in young *N. benthamiana* and tomato plants. At a concentration of 24 μM, NCR044.1 was sprayed onto the leaves of 4-week-old *N. benthamiana* and 15-day-old tomato plants and allowed to dry. The Kruskal-Wallis test showed significant differences in disease severity between different treatment groups of *N. benthamiana* (K-W_statistic_ = 22.75, df = 2, *p* < 0.0001) and tomato plants (K-W_statistic_ = 29.06, df = 2, *p* < 0.0001). *N. benthamiana* and tomato plants sprayed with the peptide had significantly reduced disease symptoms at 48 h (pairwise Wilcoxson test, corrected *p* < 0.01, Fig. 9a, b) and 60 h (pairwise Wilcoxson test, corrected *p* < 0.01, Fig. 9c, d) post-inoculation, respectively, compared to control plants sprayed with pathogen only. The photosynthetic efficiency (Fv/Fm) of the control plants (no peptide) exposed to pathogen was significantly lower compared to plants sprayed with NCR044.1 prior to pathogen exposure. In an effort to determine if defensin internalization/localization observed in fungi also occurred with plant cells, we applied a droplet of 24 μM DyLight550-NCR044.1 to the leaf surface of *N. benthamiana* and imaged it by confocal microscopy. Our results indicated that DyLight550-NCR044.1 remained on the plant surface, concentrating at anti-clinal walls, with no evidence of internalization within the leaf epidermal cells (Supplemental Fig. 6a-i). This data indicated significant potential for the use of NCR044.1 as a spray-on peptide fungicide for the control of gray mold disease in vegetables, flowers and other economically important crops.

**Figure 9.**
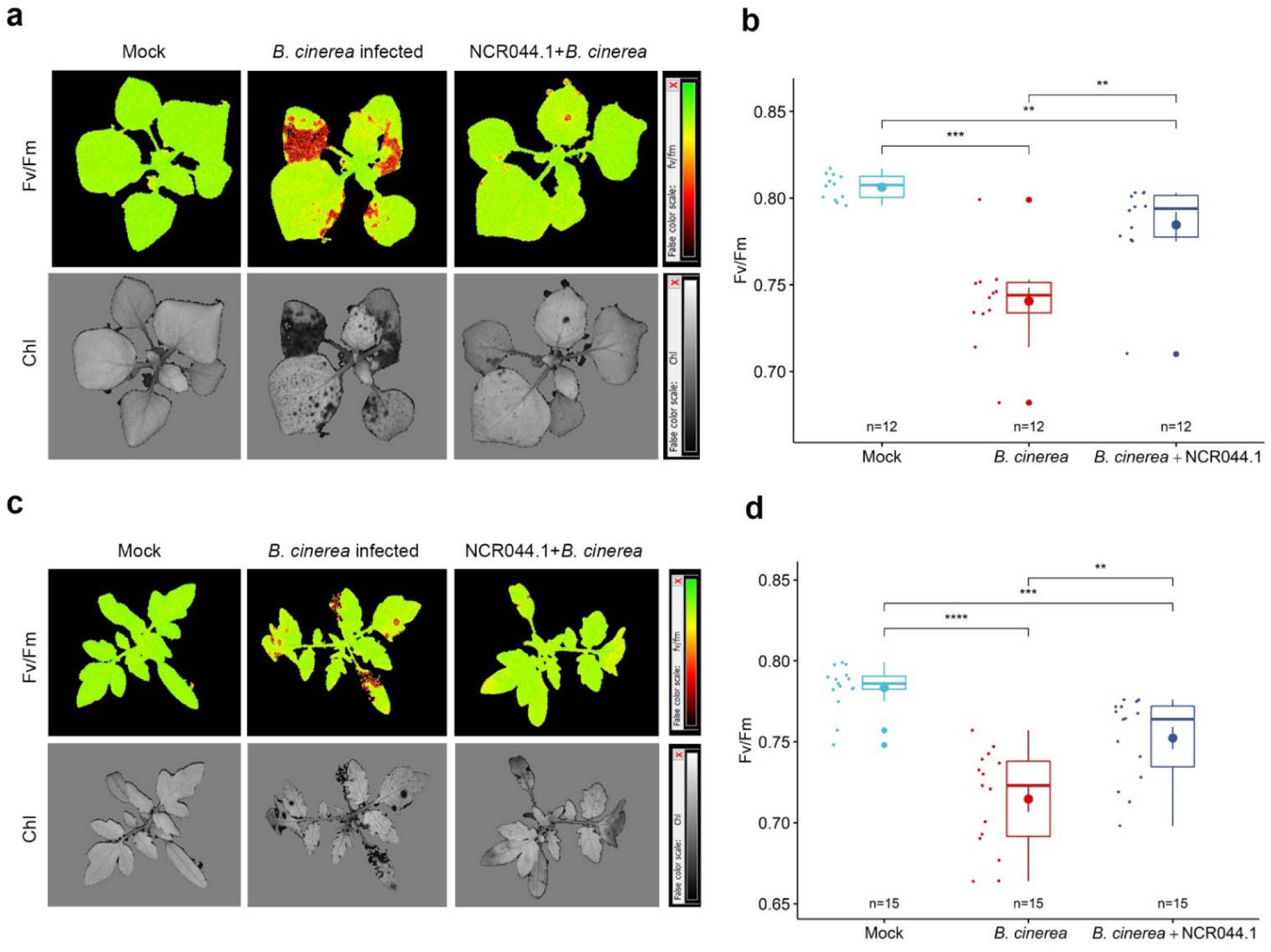
Spray application of NCR044.1 confers gray mold resistance in *N. benthamiana* and tomato plants. Four-week-old *Nicotiana benthamiana* (**a**) and two-week-old *Solanum lycopersicum* L. cv. *Mountain Spring* plants (**c**) sprayed with either 2 mL water or NCR044.1 (24 μM) prior to exposure with a spray containing 1 mL of a 5 × 10^4^ *B. cinerea* fungal spore suspension. Maximum quantum yield of PSII photochemistry (Fv/Fm) and the chlorophyll (Chl) index were measured at 48 h and 60 h, respectively, after fungal exposure. The false color gradient scale next to the figure indicates the efficiency of photosynthesis. Red (Fv/Fm) or black (Chl) colors represent a low efficiency of photosynthesis, characteristic of stress caused by the interaction between the fungus and plant. Green (Fv/Fm) or grey (Chl) colors indicate a higher efficiency of photosynthesis. NCR044.1 significantly protected tobacco and tomato plants from gray mold disease caused by *B. cinerea*. The calculated photosynthetic quantum yield (Fv/Fm) of tobacco (**b**) and tomato (**d**) plants. Each colored boxplot represents the 25^th^ and 75^th^ percentiles (box) and the whisker represents 1.5 times the interquartile range (IQR) from the 25th and 75th percentiles. The horizontal line in the box plot represents the median with the mean ± SEM denoted by a dot. Outliers are indicated by large dots outside 1.5*IQR above the first and third quartile. The individual measurements of Fv/Fm are indicated on the left of each boxplot and represent the variances of the samples. The number (n = 12) of plants tested is indicated below each group in the boxplot and is from at least two independent experiments for tobacco. Similar results were observed in a third independent experiment, except the data was collected 60 h after fungal exposure. The number (n = 15) of plants tested is indicated below each group in the boxplot and is from at least three independent experiments for tomato. Asterisks represent significant differences between different groups (*P<0.05, **P<0.01 ***P<0.001, ****≤0.0001, pairwise Wilcoxson test with Holm correction, Kruskal–Wallis for multiple groups).

## Discussion

*M. truncatula* expresses ∼639 NCR peptides during the establishment of successful symbiosis with *Sinorhizobium meliloti*^4^. These peptides are thought to be involved primarily in causing differentiation of the rhizobacteria into bacteroides. However, a subset of these peptides with high cationicity exhibit antimicrobial activity *in vitro* and *in planta*^31^. Several NCR peptides have now been shown to exhibit bactericidal activity against various Gram-negative and Gram-positive bacteria *in vitro*^3, 32^. Among the hundreds of NCR peptides expressed are two NCR044 peptides that share 83% sequence identity and high cationicity. In this study, we report the structure, antifungal activity, and MOA of one of these peptides, NCR044.1. In addition, we demonstrate its potential for control of pre- and postharvest fungal diseases.

Pre- and postharvest fungal diseases cause substantial losses in the yields of many important crops. These diseases are currently mitigated primarily through application of chemical fungicides multiple times. Unfortunately, such chemical fungicides pose considerable health and environmental risks and become ineffective over time due to the evolution of resistance. Alternative means for control of fungal infections are peptide-based biofungicides applied topically to plants^33^. NCR044.1 exhibits potent broad-spectrum antifungal activity *in vitro* against four economically important fungal pathogens tested in this study. When applied on the surface of lettuce leaves and rose petals, it significantly reduces gray mold disease lesions caused by *B. cinerea* infection. Moreover, spray application of this peptide controls gray mold disease symptoms in young tomato and *N. benthamiana* plants. These results clearly demonstrate the potential value of NCR044.1 as a spray-on peptide-based biofungicide in agriculture. In contrast to chemical fungicides, peptide-based “green chemistry” fungicides are sustainable, inexpensive, and ideal for protection of the environment and consumer health^33^.

In the inverted repeat-lacking clade legumes such as *M. truncatula*, defensins and defensin-like NCR peptides have co-evolved. Defensins are one of the first lines of defense against harmful pathogens, whereas defensin-like NCR peptides are effectors inducing differentiation of rhizobia into nitrogen-fixing bacteroids^5^. Antifungal plant defensins with four disulfide bonds are diverse in their amino acid sequences but their three-dimensional structures with a highly conserved cysteine-stabilized α/β motif are strikingly similar. To our knowledge, no three-dimensional structure of an NCR peptide has been determined to date. The structure of NCR044.1 reported here appears to be novel, dominated by largely disordered regions. It remains to be determined if this disordered structure adopts an ordered secondary structure in membrane-mimetic solutions and contributes to the biochemical function and/or stability of this peptide. Several hundred NCR peptides expressed in the nodules of *M. truncatula* also exhibit significant variation in their amino acid sequences, and therefore, they too may differ substantially in their structures and antifungal modes of action^36^.

A common characteristic of antifungal peptides is their ability to permeabilize the plasma membrane of fungal pathogens. In plant defensins, membrane permeabilization is also an important early step in their modes of action^34, 37^. Recently, the peptides NCR192, NCR247, and NCR335 have been reported to disrupt the plasma membrane of *C. albicans*^8^. In the mechanistic studies employing NCR044.1 here, we determined that NCR044.1 quickly binds to the cell walls of *B. cinerea* conidia and germlings and subsequently breaches the plasma membrane of these cells near the attachment sites. Within 30 min after peptide exposure, SYTOX Green uptake was observed at a dose-dependent rate. Thus, membrane permeabilization is also an early stage in fungal cell death induced by NCR044.1. This was further supported by our observation that as cytoplasmic loading of DyLight550-labeled NCR044.1 peptide occurred, it coincided with concomitant loss of turgor. However, conidial cells exposed to NCR044.1 for a period of 1 h could subsequently germinate, suggesting membrane permeabilization alone was insufficient to induce cell death or inhibit conidial germination.

Since several plant defensins bind to plasma membrane resident bioactive phosphoinositides or related phospholipids to oligomerize and induce membrane permeabilization^35^, we tested whether NCR044.1 also bound to specific phospholipids. Like the plant defensins NaD1^38^ and MtDef5^39^, NCR044.1 bound to several phospholipids in the phospholipid-protein overlay assay. Of note, it bound more effectively to PI mono/diphosphates, in particular, PI(3,5)P_2_. Further studies are needed to decipher the mechanism by which membrane permeabilization is induced by NCR044.1 and to understand the role phospholipid binding plays in peptide-induced membrane permeabilization of fungal pathogens.

Inhibition of mitochondrial respiration leads to the generation of ROS that play an important role in fungal cell death^34, 35^. Using the dye H_2_DCF-DA, ROS generation was detected in NCR044.1 challenged germlings, but not the conidia, of *B. cinerea*. This observation is consistent with the overall reduced peptide uptake in quiescent spores relative to germ tubes and the requirement for overnight incubation (Fig. 5a). However, further studies are needed to unambiguously establish NCR044.1-induced ROS generation as a cause of cell death. In particular, it will be important to obtain evidence for the induction of the markers of ROS-mediated oxidative damage, such as protein carbonylation and DNA laddering, that lead to apoptotic cell death.

Internalization of NCR044.1 into the conidial and germling cells of *B. cinerea* has been studied in relation to its antifungal activity. While NCR044.1 internalization by conidia was slow, it was internalized rapidly by germlings of this pathogen. Our co-localization experiments with FM4-64, an amphiphilic styryl dye that serves as a marker for endocytosis and vesicle trafficking, provided important clues about NCR044.1 initial entry into fungal cells. The earliest signs of internalization of DyLight550-labeled NCR044.1 were often indicated by a single patch near the germling tip (Fig. 6g, Fig. 6j T=0) and smaller foci just beneath the cell wall (Fig. 6g-i) which co-labeled with FM6-64. This pattern was similar to that reported for *M. truncatula* defensin MtDef5 in *Neurospora crassa*^39^ and consistent with the apical vesicle accumulation and sites of endocytosis observed with FM4-64 and AM4-64 studies in a number of diverse fungi, including *B. cinerea*^40, 41^. Once internalized, the peptide appeared to diffuse through the cytoplasm and NCR044.1 gradients formed (Fig. 6j) that coincided with origins of initial cell surface foci but no longer matched the classical membrane/vesicle labeled pattern of FM4-64 (Fig. 6d-i). Once cytoplasmic, the peptide localized to nucleoli where it likely bound ribosomal RNA and/or ribosomal proteins and inhibited protein translation. Interestingly, NCR247 has also been shown previously to interact with ribosomal proteins in the symbiont *S. meliloti*^12^. While the exact mechanism for nucleolar binding is unknown, electrostatic maps of ribosomal subunits show large areas of negative potential^42^ and this property significantly influences interactions with positively charged proteins^43^, and conceivably, NCR044.1 with a net charge of +9. It will be important to determine if other antifungal NCR peptides from IRLC legumes also target nucleoli in fungal cells. Interaction with ribosomes and inhibition of translation is likely to be a shared mechanism used by some antifungal NCR peptides^12^. However, the possibility remains that these peptides have multiple intracellular targets to induce fungal cell death.

Based on the results reported here, we propose a multistep model for the antifungal action of NCR044.1 against *B. cinerea* (Fig. 10). The first step involves binding of the peptide to the cell wall followed by binding and internalization at putative sites of endocytosis and then disruption of the plasma membrane via interaction with membrane-resident phospholipids (after spore germination). The peptide eventually gains entry into the cytoplasm, inducing the formation of ROS and binding to nucleoli where it likely forms complexes with ribosomal RNA and/or ribosomal proteins to affect translation. Our data show that NCR044.1 interacts with several major and diverse cellular components, perhaps permitted by its highly dynamic structure as determined by NMR, thus allowing broad effectiveness against fungal pathogens. These new insights necessitate further studies to better understand the relative contribution of each step to the antifungal action of NCR044.1. Hundreds of NCR peptides expressed in nodules of certain legumes offer highly diverse antifungal peptide sequence space and this work on NCR044.1 peptide structure and its MOA will help in the design of more effective antifungal peptides for future use in agriculture.

**Figure 10.**
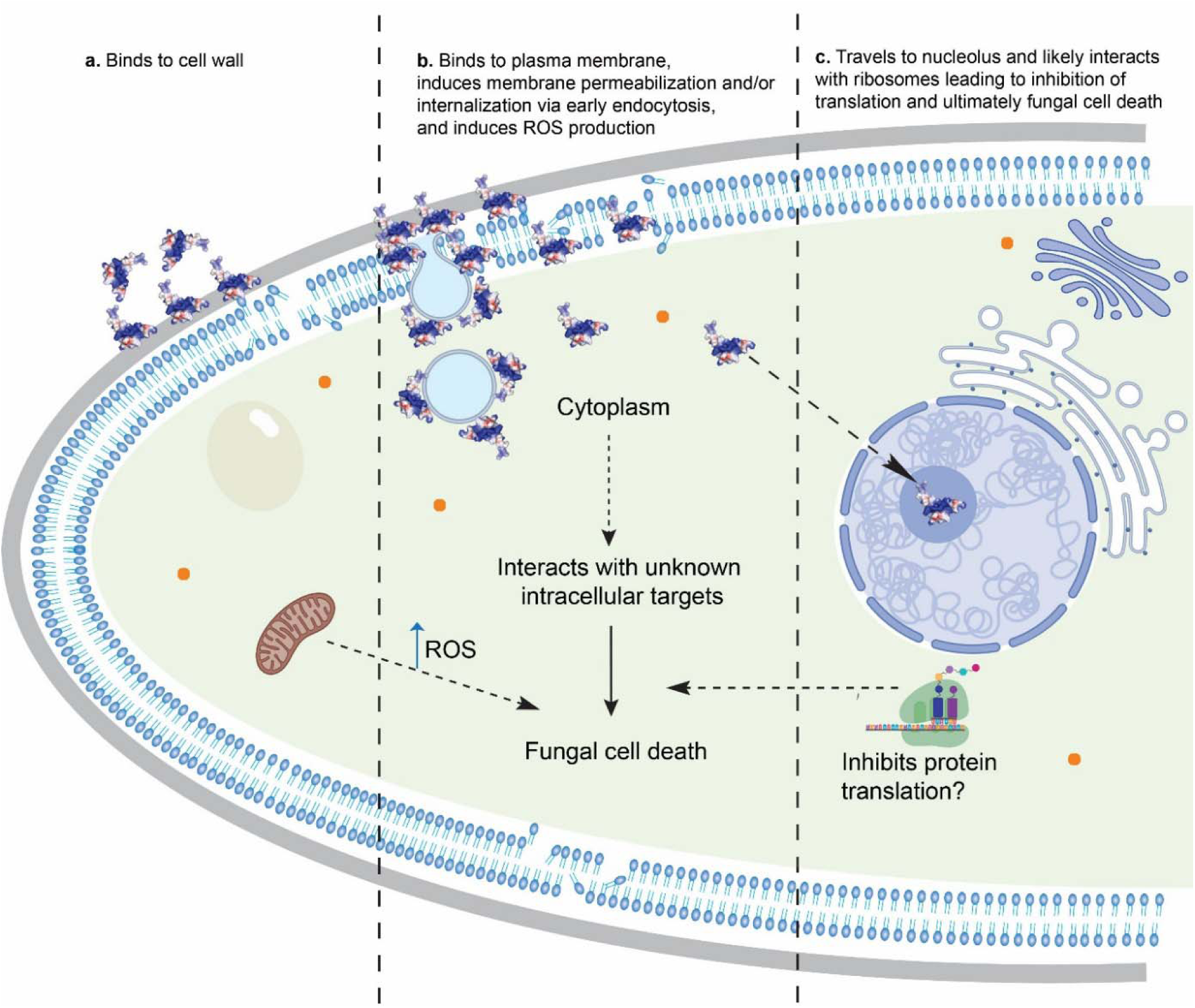
Proposed multi-step model for the antifungal action of NCR044.1 against *B. cinerea*. **a.** NCR044.1 binds to cell wall **b.** NCR044.1 binds to plasma membrane, induces membrane permeabilization and loss of turgor pressure, is internalized and induces ROS production **c.** NCR044.1 diffuses through the cytoplasm to nucleoli and likely interacts with ribosomes leading to inhibition of translation and ultimately fungal cell death.

## Methods

### Fungal cultures and spore suspensions

The fungal strains *Fusarium graminearum* PH-1, *F. virguliforme* NRL 22292 (Mont-1), *F. oxysporum* f. sp*. Cubense*, and *Botrytis cinerea* T-4 were each cultured in their respective normal growth medium shown in Supplementary Table 2. Fungal spores were harvested by washing the fungal growth plates with sterile water. The spore suspension was filtered through two layers of Miracloth, centrifuged at 13,000 rpm for 1 min, washed, and resuspended in low-salt Synthetic Fungal Medium (SFM)^44^. The spore suspension of each fungal pathogen was adjusted to the target concentration using a hemocytometer.

### Recombinant expression and purification of NCR044.1

The codon-optimized gene encoding NCR044.1 was synthesized by GenScript (Piscataway, NJ). The NCR044.1 gene was expressed in *P. pastoris* X33 using either a SacI or PmeI-linearized pPICZ α-A integration vector (Invitrogen, Carlsbad, CA) at XhoI and XbaI restriction sites in-frame with the α-mating factor secretion signal sequence containing KEX2 without the Glu-Ala repeats. The peptide product was purified from the culture medium as described previously^45^. The lyophilized peptide was dissolved in nuclease-free water and its concentration determined on a Nanodrop 2000c (Thermo Scientific, Waltham, MA) using A280 with the molar extinction coefficient (3230 M^-1^cm^-1^) and molecular weight (4.3 kDa) of NCR044.1. The purity and mass of NCR044.1 were verified by sodium dodecyl sulfate-polyacrylamide gel electrophoresis (SDS-PAGE) and direct infusion-mass spectrometry, respectively.

### ^15^N and ^13^C isotopic labeling of NCR044.1 for NMR structural analysis

The ^13^C- and ^15^N-labeled peptide was prepared according to the protocol as described previously^46^ with minor modifications. The labeled NCR044.1 was purified from the *P. pastoris* growth medium following a protocol similar to the one used for the unlabeled peptide^45^.

### NMR spectroscopy of NCR044.1

All NMR data were collected at 20°C on a double-labeled (^13^C, ^15^N) sample of NCR044.1 (0.7 mM, 20 mM sodium acetate, 50 mM NaCl, pH 5.3) using Agilent (Inova-600 and −750) or Bruker (Advent-750) spectrometers equipped with an HCN-probe and pulse field gradients. The ^1^H, ^13^C, and ^15^N chemical shifts of the backbone and side chain resonances were assigned from the analysis of two-dimensional ^1^H-^15^N HSQC, ^1^H-^13^C HSQC, HBCBCGCDHD, and HBCBCGCDCHE spectra and three-dimensional HNCACB, CBCA(CO)NH, HCC-TOCSY-NNH, CC-TOCSY-NNH, and HNCO spectra. The chemical shift assignments were assisted by the analysis of three-dimensional ^15^N-edited NOESY-HSQC and ^13^C-edited NOESY-HSQC (aliphathic and aromatic) spectra, collected with a mixing time of 90 ms, that also supplied the NOE-based distance restraints required for the structure calculations. To improve the resolution around the residual water resonance and obtain additional inter- and intra-residue proton side chain restraints, a three-dimensional ^13^C-edited NOESY-HSQC (aliphathic) data set was collected by lyophilizing and resuspending the sample in 99.8% D_2_O. In preparing the latter sample, slowly exchanging amides were identified by collecting a short ^1^H-^15^N HSQC spectrum after the addition of 99.8% D_2_O (∼10 minutes later). To assess the effect of the disulfide bonds on the structure of NCR044.1, the double-labeled (^13^C, ^15^N) sample in 99.8% D_2_O was lyophilized, redissolved in 93% H_2_O/7% D_2_O, and made 5 mM with the strong reducing agent tris(2-carboxyethyl)phosphine (TCEP). Similar two- and three-dimensional NMR experiments used to assign the ^1^H, ^13^C, and ^15^N chemical shifts of oxidized NCR044.1 were used to assign these resonances in the reduced state. An overall rotational correlation time, τ_c_, was estimated for NCR044.1 in the reduced and oxidized states from the ratio of collective backbone amide ^15^N T_1_ and T_1rho_ measurements^47^. A chemical shift perturbation study^29^ was performed with a 250 μL sample of oxidized NCR044.1 (0.07 mM) by titrating four times 2 μL aliquots of phosphatidylinositol 3-phosphate diC8 (PI(3)P, 42.5 mM) (Echelon Biosciences, Salt Lake City, UT) and collecting a ^1^H-^15^N HSQC after each addition. The molar ratio of PI(3)P:NCR044.1 at each titration point was 4.7, 9.4, 14.1, and 18.8. All NMR data were processed using Felix2007 (MSI, San Diego, CA) software and analyzed with the program Sparky (v3.115). The ^1^H, ^13^C, and ^15^N chemical shifts were referenced using indirect methods (DSS = 0 ppm) and deposited into the BioMagResBank database (www.bmrb.wisc.edu) under the accession number BMRB-30660 (oxidized NCR044.1).

### NMR solution structure calculations for oxidized NCR044.1

The ^1^H, ^13^C, and ^15^N chemical shift assignments and peak-picked NOESY data were used as initial experimental inputs in iterative structure calculations with the program CYANA (v 2.1)^48^. The assigned chemical shifts were also the primary basis for the introduction of 38 dihedral Psi (ψ) and Phi (φ) angle restraints identified with the program TALOS+ using the online webserver (https://spin.niddk.nih.gov/bax/nmrserver/talos/)^49^. Six restraints between the side chain sulfur atoms of C9-C30 and C15-C25 (2.0-2.1 Å, 3.0-3.1 Å, and 3.0-3.1 Å for the Sγ-Sγ, Sγ-Cβ, and Cβ-Sγ distances, respectively)^21^ were introduced on the basis of high resolution mass spectrometry analysis. Four hydrogen bond restraints (1.8 – 2.0 Å and 2.7 – 3.0 Å for the NH–O and N-O distances, respectively) were introduced on the basis of proximity in early structure calculations and the observation of slowly exchanging amides in a deuterium exchange experiment. The final ensemble of 20 CYANA derived structures were then refined with explicit water with CNS (version 1.1) using the PARAM19 force field and force constants of 500, 500, and 1000 kcal for the NOE, hydrogen bond, and dihedral restraints, respectively. For these water refinement calculations the upper boundary of the CYANA distance restraints was left unchanged and the lower boundary set to the van der Waals limit. Structural quality was assessed using the online Protein Structure Validation Suite (PSVS, v1.3)^50^. Complete structural statistics are summarized in Table 1. The atomic coordinates for the final ensemble of 19 structures have been deposited in the Research Collaboratory for Structural Bioinformatics (RSCB) under PDB code 6U6G.

### Antifungal activity assay

The antifungal activity of NCR044.1 against fungal pathogens was determined spectrophotometrically using the 96-well plate assay as described previously^51, 52^. The quantitative fungal growth inhibition was determined by measuring the absorbance at 595 nm using a (Tecan Infinite M200 ProTecan Systems Inc., San Jose, CA) microplate reader at 48 h. IC_50_ and IC_90_ values were calculated with GraphPad software (version 8, San Diego, CA). Fungal cell viability/cell killing was determined using the resazurin cell viability assay as described previously^52, 53^.

### SYTOX^TM^ Green (SG) membrane permeabilization assay

The effect of NCR044.1 on the membrane integrity of *B. cinerea* germlings was determined using a modified SG uptake quantification assay^45^. Membrane permeabilization in *B. cinerea* germlings exposed to different concentrations of NCR044.1 was measured every 30 min for 2 h. SG uptake was quantified by measuring the fluorescence in a SpectraMax® M3 spectrophotometer (exc: 488; cutoff: 530; emi: 540 nm). Membrane permeabilization of *B. cinerea* conidia (50 μl, 10^5^ spores/ml) and germlings was measured after 15 min of exposure to NCR044.1 (50 μL, 3 μM) using fluorescence confocal microscopy (Leica SP8-X) with an excitation and emission wavelength of 488 and 508-544 nm, respectively.

### Quantification of intracellular reactive oxygen species (ROS)

ROS production in *B. cinerea* germlings exposed to different concentrations NCR044.1 was measured every 30 min for 2 h using the ROS indicator dye 2′,7′-dichlorodihydrofluorescein diacetate (H_2_DCF-DA, Invitrogen, Carlsbad, CA). ROS levels were quantified by measuring the fluorescence in a SpectraMax® M3 spectrophotometer microplate reader (exc: 495; cutoff: 515; emi: 529 nm). Intracellular ROS production in *B. cinerea* conidia and germlings exposed to 3 μM NCR044.1 was also imaged using fluorescence confocal microscopy (Leica SP8-X)^39^ with an excitation and emission wavelength of 480 and 510-566 nm, respectively.

### NCR044.1 uptake into fungal cells

NCR044.1 was labeled with DyLight550 amine reactive fluorescent dye following the manufacturer’s instructions (Thermo Scientific, Waltham, MA). Confocal microscopy on a Leica SP8-X was performed to monitor uptake of the fluorescently labeled NCR044.1 peptide as described previously^54^ with minor modifications. Briefly, *B. cinerea* conidia (50 μl, 10^5^ spores/ml) were treated with DyLight550-labeled NCR044.1 (50 μL, 1.5 and 3 μM), incubated at room temperature overnight, and the fluorescence measured with excitation and emission wavelengths of 550 and 569-611 nm, respectively. To monitor internalization of DyLight550-NCR044.1 into *B. cinerea* germlings, 3D time-lapse confocal microscopy data were collected every 3 min for 1 h. Co-localization experiments were performed to visualize the intracellular distribution of DyLight550-NCR044.1 and the membrane-selective dye FM4-64 using an excitation wavelength of 550 nm for both dyes with 560-600 and 680-800 nm emission wavelengths, respectively. To determine if NCR044.1 targeted the nuclear DNA or nucleolus region of *B. cinerea*, germlings were treated with 3 μM of DyLight500-NCR044.1 in the presence of the DNA specific staining dye 4′,6-diamidino-2-phenylindole (DAPI, final concentration: 0.1 μg/mL) or a RNA selective nucleolus specific staining dye (Nucleolus Bright Green, final concentration: 1 μmol/L, Dojindo Molecular Technologies, Japan). Co-localization was visualized using a ZEISS Elyra 7 super-resolution microscope (Carl Zeiss Microscopy, Jena, Germany) in lattice structured illumination microscopy mode at excitation and emission wavelengths of 561 and 590 nm (DyLight550), 405 and 440 nm (DAPI), and 488 and 515 nm (Nucleolus Bright Green), respectively. Single- or time-lapse z-stacks were acquired with a C-Apochromat 63X water immersion objective lens (Numerical aperture = 1.2, 63 nm x-y and 371 nm z pixel size in leap mode) taking 15 phases per image with a 30 ms exposure.

### Fluorescence Redistribution After Photobleaching (FRAP) experimental setup and analysis

The FRAP experiments were performed on a Leica SP8-X microscope (FRAP Wizard Module, LAS X Version 3.1.5.16.16308) using the HC PlanApochromat 63X water immersion objective lens (Numerical aperture = 1.2) with the pinhole set to 1 Airy unit and zoom set to 1.77. Images were acquired for the selected cellular regions (germling cytoplasm, germling nucleoli, germling cell wall, and spore cell wall) after 1 h incubation with peptide under the following identical imaging and FRAP conditions: 10 pre-bleach and 700 post-bleach frames (512 x 64 pixels, pixel size 204 nm) captured at 173 ms intervals without averaging using 550 nm excitation and 560-625 nm emission wavelengths with a PMT detector and a transmitted light channel. The power of the 550 nm laser was set at 3.5% during imaging to prevent the detection of bleaching for the duration of each FRAP experiment (∼3 min). A 1.52 μm^2^ circular bleach region was applied with 100% laser excitation power (550 nm wavelength) for 3.5 s immediately following pre-bleach and prior to post-bleach imaging. FRAP analysis was performed on 24 datasets for each subcellular region showing no evidence of sample drift during acquisition. A single exponential curve fitting function was applied post-acquisition to extract the half-maximum tau (τ½), mobile fraction, and immobile fraction for each condition.

### Peptide distribution on the plant surface

The distribution of DyLight550-NCR044.1 on tobacco leaf surfaces was characterized by incubating leaves with 24 uM of labeled peptide for 1 h, air drying, and imaging on a Leica SP8-X confocal microscope equipped with an HC PlanApochromat 63X water immersion objective lens (Numerical aperture = 1.2) with the pinhole set to 1 Airy unit. Simultaneous excitation with 405 and 560 nm laser wavelengths was used to visualize cell wall auto-(426 – 485 nm) and DyLight550-(560 – 600 nm) fluorescence using photo-multiplier tube (PMT) detection of a transmitted light channel. Three-dimensional volumes were created using a 1 μm z-interval for each treatment and rendered with a 3D maximum intensity projection.

### NCR peptide-phospholipid interactions

NCR044.1-phospholipid interactions were determined using PIP Strips^TM^ (Echelon Biosciences, Salt Lake City, USA) as described previously^54^ with minor modifications. The binding of NCR044.1 to the phospholipids on the strips was detected using a polyclonal antibody generated in rabbits with custom-synthesized NCR044.1 (Biomatik Corp., Cambridge, ON, CDN). Binding of NCR044.1 to PI4P and PI(3,5)P_2_ was determined using the PolyPIPosomes (Echelon Biosciences, Salt Lake City, USA) assay as described^38^. Samples were reduced with 100 mM dithiothreitol (DTT), separated on a 4-20% (Bio-Rad, TGX™ precast gels) SDS-PAGE gel, and then stained with Instant Blue (Expedeon, Cambridge, UK).

### Semi-in *planta* antifungal activity of NCR044.1 against *B. cinerea*

Detached rose petal and lettuce leaf infection assays were performed as described previously^55^ with minor modifications. Briefly, store-bought rose petals or ∼2 x 2 cm cut pieces of iceberg lettuce were placed in petri dishes. A 10 μl aliquot of NCR044.1 was spotted onto the plant samples at different peptide concentrations and inoculated with 10 μL of *B. cinerea* spores (10^5^ spores/mL suspended in 2X SFM buffer). The petri dishes were inserted into Ziploc WeatherShield plastic boxes containing a wet paper towel to maintain a high humidity. Following incubation at room temperature for 48 h the leaves were photographed and the lesion size was determined using ImageJ software.

### Evaluation of NCR044.1 as sprayable peptide for protection against gray mold disease

The tomato (*Solanum lycopersicum* cv. Mountain Spring) and tobacco (*Nicotiana benthamiana*) plants were grown in a controlled environment growth chamber for two and four weeks, respectively, under a 16/8 h and 12/12 h light/dark cycle, respectively. Whole plants were then sprayed with water or NCR044.1 (24 μM), allowed to dry for 1 h, sprayed with a 5 × 10^4^ *B. cinerea* fungal spore suspension prepared in half-strength SFM medium, and placed in a highly humid environment. After 48 h, high-resolution fluorescence images were taken using CropReporter (PhenoVation, Wageningen, Netherlands). The quantification of plant health/stress was carried out using the calculated F_V_/F_M_ (maximum quantum yield of photosystem II) images of the CropReporter, showing the efficiency of photosynthesis in false colors.

### Statistical Analysis

Statistical analysis was performed in the R environment (version 3.5.2) using RStudio (version 1.1.463). The normality and homogeneity of variance was evaluated using a Shapiro-Wilk test and F-test/Levene’s test, respectively. A parametric unpaired Student’s *t* test was used to compare the significant difference between two groups of normally distributed data. The non-parametric Kruskal-Wallis test was performed on non-normally distributed data to compare significant differences of more than two groups. A pairwise Wilcoxson test with an Holm-Bonferroni p-value correction was applied to find significant differences between groups.

## Acknowledgments

The generous support from TechAccel for part of the research reported here is acknowledged. The structure for NCR044.1 was solved using resources at the W.R. Wiley Environmental Molecular Sciences Laboratory, a national scientific user facility sponsored by U.S. Department of Energy’s Office of Biological and Environmental Research (BER) program located at Pacific Northwest National Laboratory (PNNL). Battelle operates PNNL for the U.S. Department of Energy. We also acknowledge imaging support from the Advanced Bioimaging Laboratory at the Danforth Plant Science Center and usage of the Leica SPX-8 acquired through an NSF Major Research Instrumentation grant (DBI-1337680).

## Author Contributions

S.V., K.C., and D.S. conceived the research. S.V., K.C., H.L., G.B., and D.S. designed the research. S.V., H.L., J.S., and G.B. conducted the experiments. S.V., K.C., H.L., J.S., G.B., and D.S. analyzed and interpreted the results. S.V., K.C., G.B., and D.S. wrote the manuscript with help from H.L.

