## Supplementary Data for "Antifungal symbiotic peptide NCR044.1 exhibits unique structure and multi-faceted mechanisms of action that confer plant protection"

Siva L. S. Velivelli *et al*.

|  | **C9** | **C15** | **C25** | **C30** |
| --- | --- | --- | --- | --- |
| **Oxidized** | 45.6 | 38.0 | 40.3 | 39.5 |
| **Reduced** | 27.9 | 27.7 | 27.9 | 28.0 |
| **Δ^13^C^β^** | 17.7 | 10.3 | 12.4 | 11.5 |

**Supplementary Table 1.** Chemical shifts (ppm) for the ^13^C^β^ carbons of the four cysteine residues in NCR044.1 in the oxidized and reduced states.

| PDB ID | 6U6G |
| --- | --- |
| BMRB ID | 30660 |
| **Restraints for Structure Calculations** |  |
| Total NOEs | 179 |
| Intraresidue NOEs | 93 |
| Sequential (i, i + 1) NOEs | 68 |
| Medium-range (i, i + j; 1 < j ≤ 4) NOEs | 6 |
| Long-range (i, i + j; j > 4) NOEs | 12 |
| Phi (Φ) angle restraints | 19 |
| Psi (Ψ) angle restraints | 19 |
| Hydrogen bonds | 4 |
| **Structure Calculations** |  |
| Number of structures calculated | 100 |
| Number of structures used in ensemble | 19 |
| **Structures with Restraint Violations** |  |
| NOE-based Distance Restraint Violations > 0.05Å | 0 |
| Dihedral Restraint Violation > 1º | 1 |
| **RMSD to Mean (Å) for PSVS Ordered Residues^b^** |  |
| Backbone N-C^^-C=O Atoms | 0.84 ± 0.31 Å |
| All Heavy Atoms | 1.78 ± 0.25 Å |
| **Ramachandran Plot Summary**  **(Richardson Lab’s Molprobity)^c^** |  |
| Most favored regions | 97.9% |
| Additionally allowed regions | 2.1% |
| Generously favored regions | 0.0% |
| Disallowed | 0.0% |
| **Global Quality Scores (Z-score (raw))^c^** |  |
| Procheck (all) | -2.48 (-0.42) |
| Procheck (Φ, Ψ) | -0.90 (-0.31) |
| MolProbity clash score | 0.15 (8.04) |

**Supplementary Table 2.** Summary of the structural statistics for the solution structures of NCR044.1^a^. ^a^All statistics are for the ensemble deposited in the Protein Data Bank. ^b^Ordered residues: S6-E14, A23-R32. ^c^Residues with secondary structure elements: D12-C15, R24-R26, Y29-V31.

| **Strains** | **Stock medium** | **Medium for spore production** | **Culture conditions** | **Reference** |
| --- | --- | --- | --- | --- |
| *F. graminearum* | PDA | CMC broth | 3-5 days, 28˚C, 180 rpm | Cappelini & Peterson (1965) |
| *F. virguliforme* | PDA | PDA | 7-14 days, 25˚C | This study |
| *F. oxysporum* | PDA | PDA |  | This study |
| *B. cinerea* | PDA | Modified V8 agar |  | This study and Lian et al., 2018 |

PDA denotes Potato Dextrose Agar, V8 denotes commercial vegetable juice V8 ((360 mL/L V8 vegetable juice, 2 g/L CaCO_3_, 15 g/L agar), and CMC denotes Carboxymethyl Cellulose Medium (15 g/L carboxymethyl cellulose, 1 g/L yeast extract, 0.5 g/L MgSO_4_.7 H_2_O, 1 g/L NH_4_NO_3_, and 1 g/L KH_2_PO_4_)

Cappellini RA, Peterson JL (1965) Macroconidium formation in submerged cultures by a non-sporulating strain of *Gibberella zeae*. Mycologia 57 (6):962-966.

Lian J, Han H, Zhao J, Li C (2018*) In-vitro* and *in-planta* *Botrytis cinerea* inoculation assays for tomato. Bio-protocol 8 (8):e2810.

**Supplementary Table 3.** Growth media and conditions for culturing the fungi used in this study.

**Supplementary Figure 1.** **Molecular weight determination of NCR044.1 by direct infusion mass spectrometry.**   **a.** The raw MS/MS data of NCR044.1. **b.** The deconvoluted mass spectrum (monoisotopic mass) of NCR044. 1 (M = 4311.28 Da).

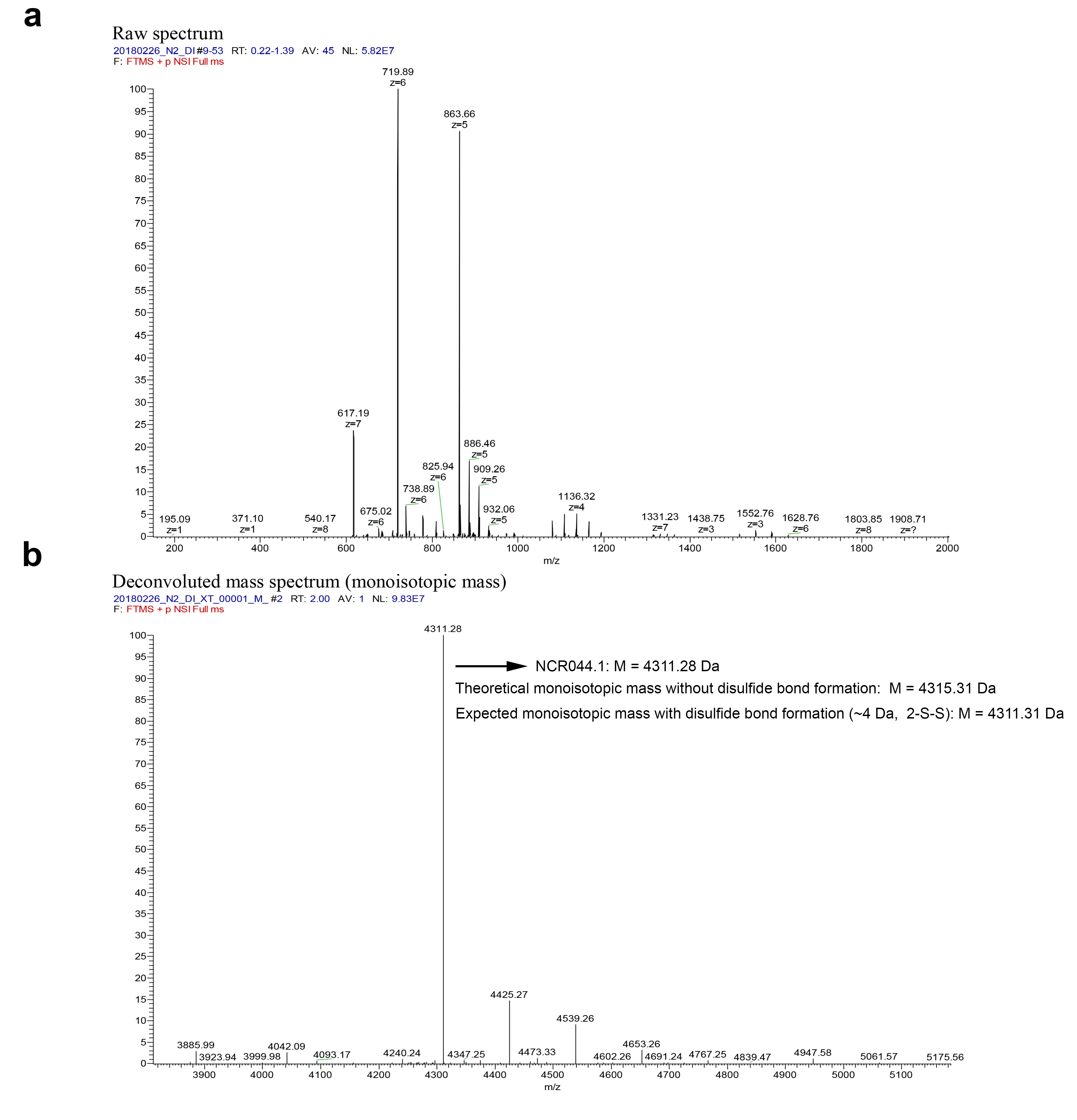

**Supplementary Figure 2**. **Effect of NCR044.1 on intracellular ROS generation in *B. cinerea.***

**a-d.** Representative confocal microscopy images showing the production of ROS (green) in H_2_DCFDA treated *B. cinerea* conidia and germlings 2 h after exposure to 3 µM NCR044.1. Fluorescence due to ROS production was only observed in *B. cinerea* germlings. (Scale bar: 10 µm). **e-h**. Representative bright field images of *B. cinerea* conidia and germlings. **i.** Real time-quantification of ROS production in *B. cinerea* germlings treated with various concentrations of NCR044.1. ROS fluorescence was observed 30 min after exposure to NCR044.1 and increased with time in a dose-dependent manner. Data are mean (large dot) ± SEM of three biological replications (n = 3, small dots).

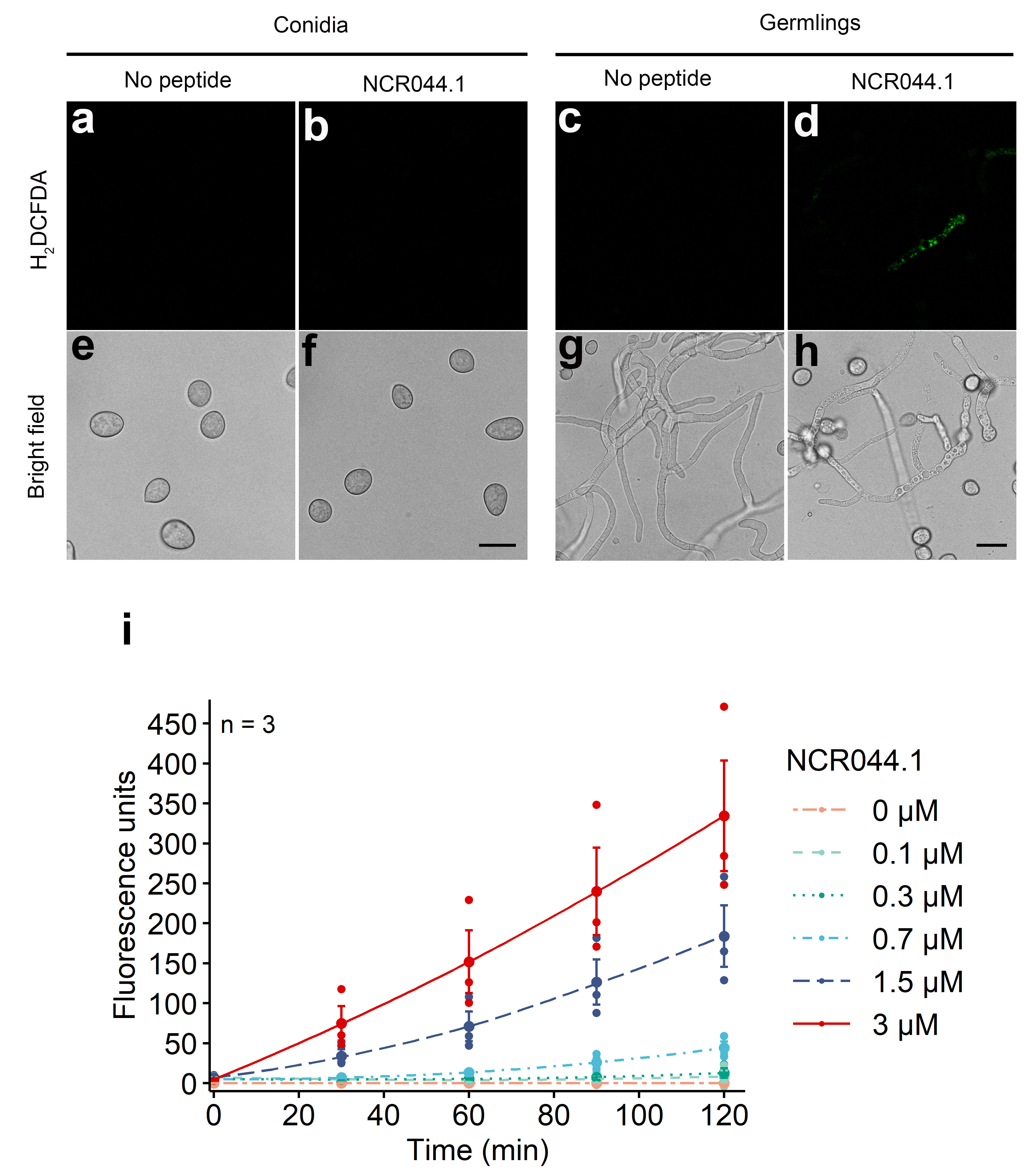

**Supplementary Figure 3. Chemical shift perturbation study with PI(3)P and map of the electrostatic potential of the solvent-accessible surface of NCR044.1.** **a.** Overlay of the ^1^H-^15^N HSQC spectra of NCR044.1 (0.07 mM) without lipid (red) and with a 18.8 molar ratio of PI(3)P:NCR044.1 (black). The circled resonances highlight amide resonances still visible at this lipid concentration. Spectra collected at 20 °C in 20 mM sodium acetate, 50 mM NaCl, pH 5.3 at a ^1^H resonance frequency of 600 MHz. **b.** *Pymol*-generated map of the electrostatic potentials on the solvent-accessible surface of NCR044.1. The map on the left was in the same orientation as structure in Figure 2d. Red and blue correspond to negatively and positively charged surfaces, respectively.

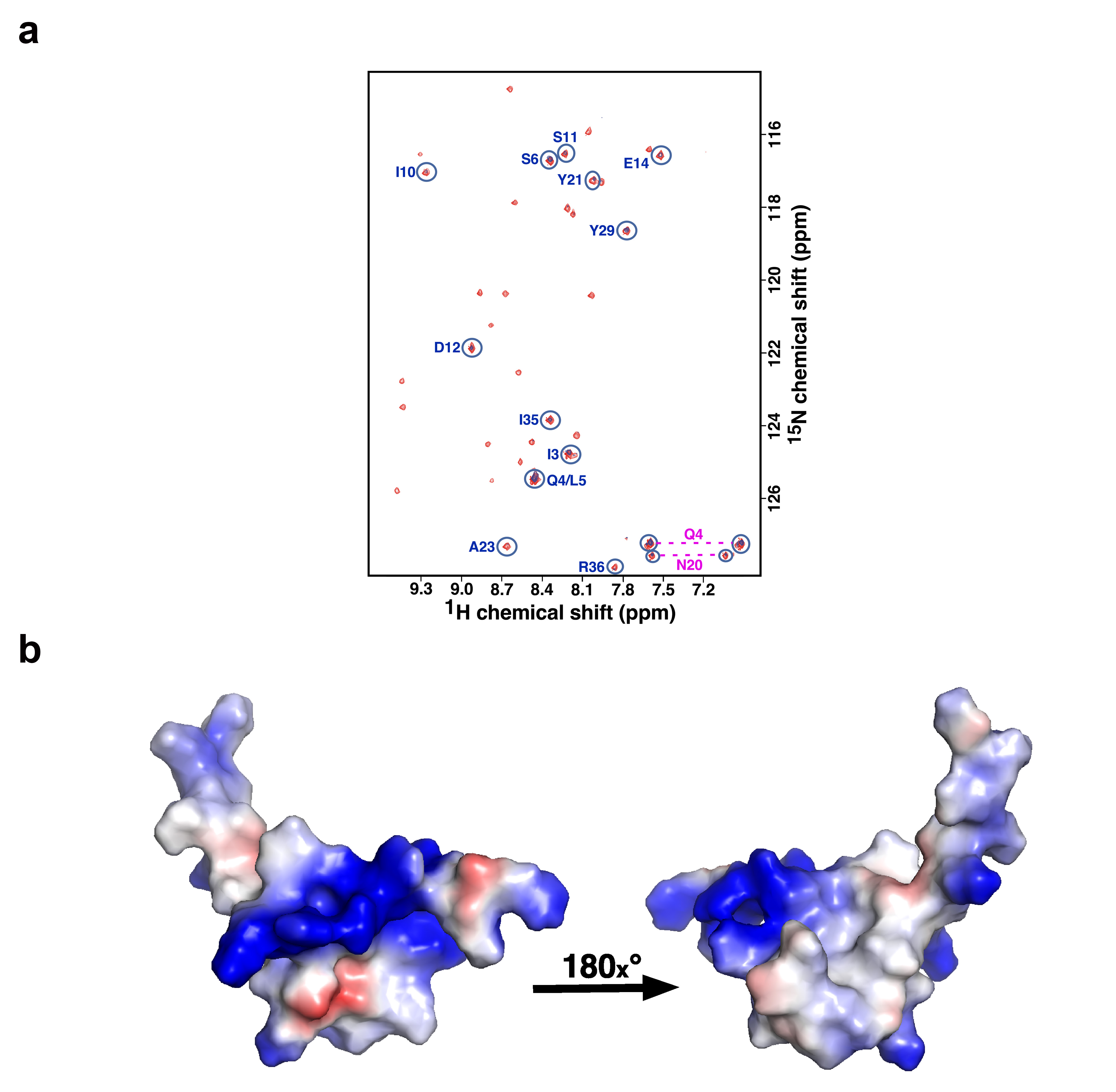

**Supplementary Figure 4**. **Exogenously applied NCR044.1 confers resistance to *B. cinerea* in lettuce leaves and rose petals**. Lettuce leaves (**a-b)** and rose petals (**c-d)** inoculated with different concentrations of NCR044.1 displayed smaller lesions 48 h after exposure to *B. cinerea*. Each colored boxplot represents the 25^th^ and 75^th^ percentiles (box) and the whisker represents 1.5 times the interquartile range (IQR) from the 25th and 75th percentiles. The horizontal line in the boxplot represents the median with the mean ± SEM denoted by a dot. Outliers are indicated by large dots outside 1.5*IQR above the first and third quartile. The individual measurements of lesion size are indicated on the left of each boxplot and represent the variances of the samples. The number (n = 36) of lettuce leaves tested is indicated below each group in the boxplot and are from at least three independent experiments. The number (n = 24) of rose petals tested is indicated below each group in the boxplot and are from at least three independent experiments. Asterisks represent significant differences between different groups (*P < 0.05, **P < 0.01 ***P < 0.001, ****≤0.0001, pairwise Wilcoxson test with Holm correction, Kruskal–Wallis for multiple groups).

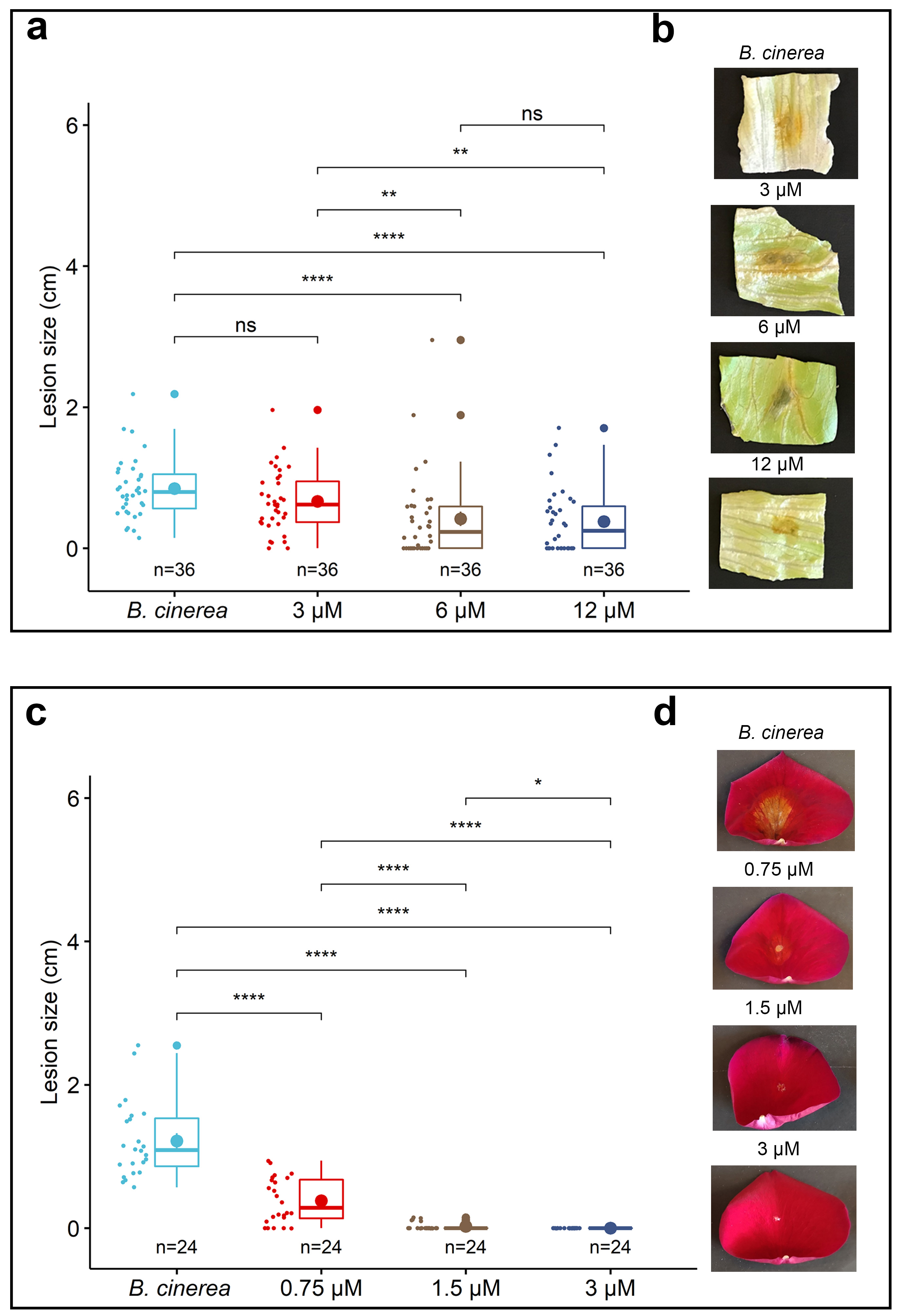

**Supplementary Figure 5**. **Localization of DyLight550-NCR044.1 in cells of the leaf of *N. benthamiana*.** **a.** Cell wall autofluorescence, **b.** DyLight550-labeled NCR044.1, **c.** Merged-maximum intensity projection, **d.** Cell wall autofluorescence - single optical slice same data set as in (**a**), **e.** DyLight550-labeled NCR044.1 excluded from the plant cytoplasm, **f.** Merged, **g.** Cell wall autofluorescence, **h.** DyLight550 only without peptide, **i.** Merged. These data show that DyLight550-labeled NCR044.1 concentrates near anticlinal walls and is not internalized inside the plant cells (Scale bar: 20 µm).

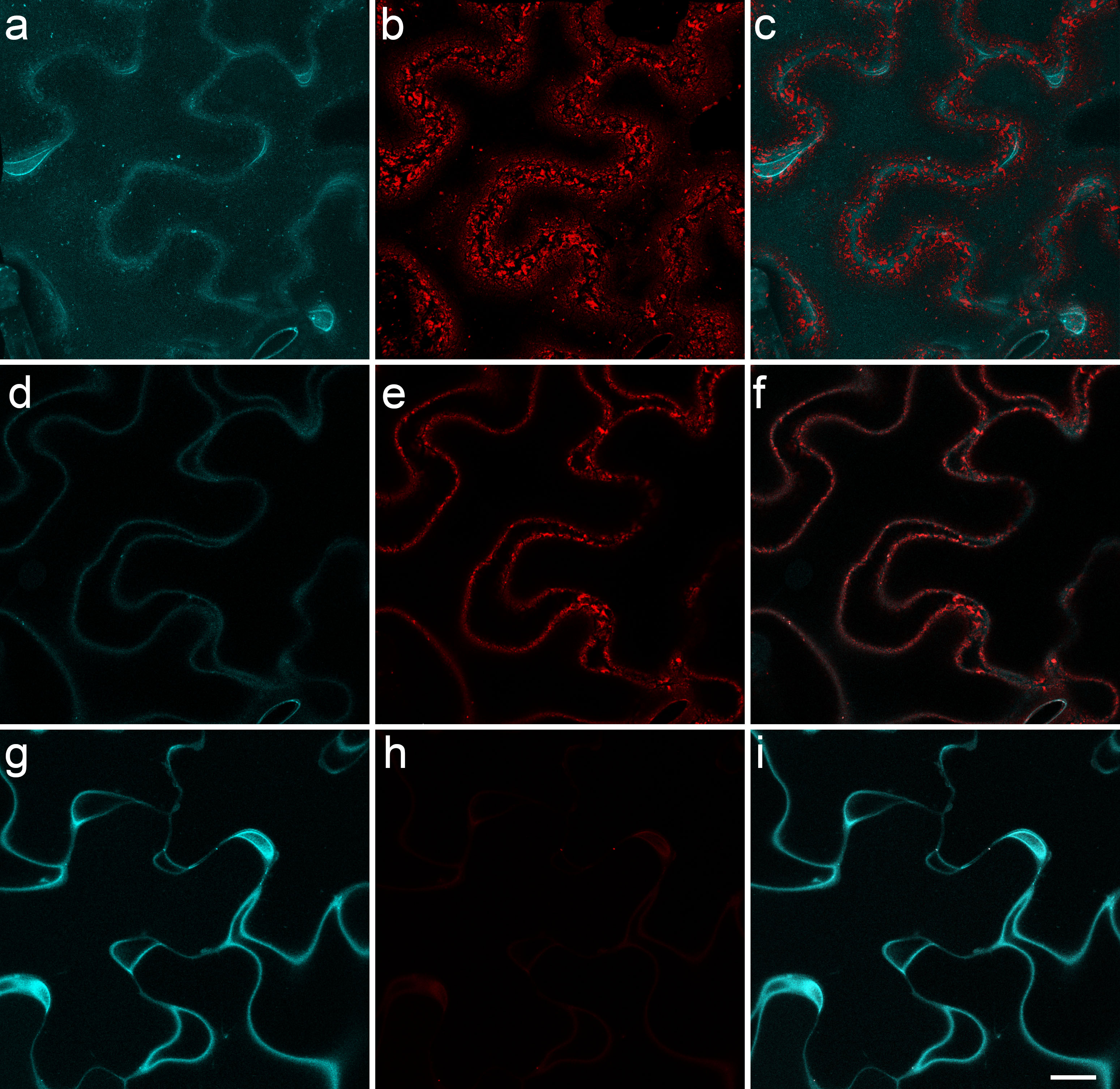
